# Estimating the timing of multiple admixture pulses during local ancestry inference

**DOI:** 10.1101/314617

**Authors:** Paloma Medina, Bryan Thornlow, Rasmus Nielsen, Russell Corbett-Detig

## Abstract

Admixture, the mixing of genetically distinct populations, is increasingly recognized as a fundamental biological process. One major goal of admixture analyses is to estimate the timing of admixture events. Whereas most methods today can only detect the most recent admixture event, here we present coalescent theory and associated software that can be used to estimate the timing of multiple admixture events in an admixed population. We extensively validate this approach and evaluate the conditions under which it can succesfully distinguish one from two-pulse admixture models. We apply our approach to real and simulated data of *Drosophila melanogaster*. We find evidence of a single very recent pulse of cosmopolitan ancestry contributing to African populations as well as evidence for more ancient admixture among genetically differentiated populations in sub-Saharan Africa. These results suggest our method can quantify complex admixture histories involving genetic material introduced by multiple discrete admixture pulses. The new method facilitates the exploration of admixture and its contribution to adaptation, ecological divergence, and speciation.

## Introduction

There is an increasing appreciation for the importance of admixture, an evolutionary process wherein genetically divergent populations encounter each other and hybridize. Admixture has shaped genetic variation within natural plant, animal, and human populations (Pool *et al*. 2012; Rieseberg *et al*. 2003; Hufford *et al*. 2013; Sankararaman *et al*. 2014; Hellenthal *et al*. 2014). If an admixture event has occurred relatively recently, we can use local ancestry inference methods (LAI) to trace the ancestry of discrete genomic segments, called ‘ancestry tracts,’ back to the ancestral populations from which they are derived (Sankararaman *et al*. 2012; Corbett-Detig and Nielsen 2017; Maples *et al*. 2013; Pool and Nielsen 2009; Price *et al*. 2009). Due to ongoing recombination within admixed populations, the lengths of these ancestry tracts are expected to be inversely related to the timing of admixture. Therefore, it is possible to estimate the timing of admixture events by inferring ancestry tract lengths (Pool and Nielsen 2009; Gravel 2012), or by evaluating the rate of decay of linkage disequilibrium (LD) among ancestry informative alleles (Moorjani *et al*. 2011; Loh *et al*. 2013).

The latter approach is based on modeling expected decay of LD among alleles that are differentiated between admixed populations. Briefly, even if there is little LD in the ancestral populations themselves, admixture will create admixture LD (ALD) within the admixed population among alleles whose frequencies are differentiated between ancestral populations (Chakraborty and Weiss 1988). The decay of ALD is expected to be approximately exponential with a rate parameter that is proportional to the timing of admixture. Two popular methods using this approach, ROLLOFF (Moorjani *et al*. 2011) and ALDER (Loh *et al*. 2013), model the decay of two-locus ALD to estimate single pulse admixture models. Admixture histories may be more complex, including multiple distinct admixture events (Gravel *et al*. 2013) and multiple ancestral populations (Pasaniuc *et al*. 2009), suggesting that some of these methods may not be suitable for deeply characterizing the admixture history of many admixed populations (Figure 1). Although, recent work suggests it is possible to detect multiple admixture pulses by modeling LD decay as a mixture of two exponential distributions (Pickrell *et al*. 2014). Additionally, it may be possible to extend LD decay approaches using three-point linkage disequilibrium to estimate the timing of two admixture pulses in populations with more complex admixture histories (Liang and Nielsen 2016). Finally, wavelet-based techniques can be used to infer the relationship between SNPs along a chromosome to estimate the time of admixture and admixture proportions (Sanderson *et al*. 2015; Pugach *et al*. 2016).

A second set of approaches uses the lengths of LA tracts across the genomes of admixed individuals to estimate the timing of admixture events (Gravel 2012; Gravel *et al*. 2013; Pool and Nielsen 2009; Ni *et al*. 2018). Here, the complexity of admixture models that can be accommodated depends on the accuracy of the estimated ancestry tract length distributions within an admixed population. However, these methods require that LAI be performed prior to estimating population admixture models. LAI necessitates *a priori* assumptions about the admixture model itself, as such assumptions are a prerequisite—implicitly or explicitly—for most LAI frameworks that have been developed to date. These assumptions have the potential to bias the outcomes of LAI and can affect admixture model selection (Pool *et al*. 2012; Sankararaman *et al*. 2008; Corbett-Detig and Nielsen 2017). Additionally, LAI methods require accurately phased chromosomes—an unlikely prospect for species with low levels of LD (Bukowicki *et al*. 2016) and for relatively ancient admixture events where the rate of phasing switch errors would be similar to the rate of transitions between ancestry types. Therefore, it is often preferable to estimate the timing of admixture events and the LA of a sample simultaneously (Corbett-Detig and Nielsen 2017). Finally, modern sequencing techniques often sequence to light coverage, thus necessitating a tool that can accommodate read pileup data rather than genotypes.

**Figure 1.**
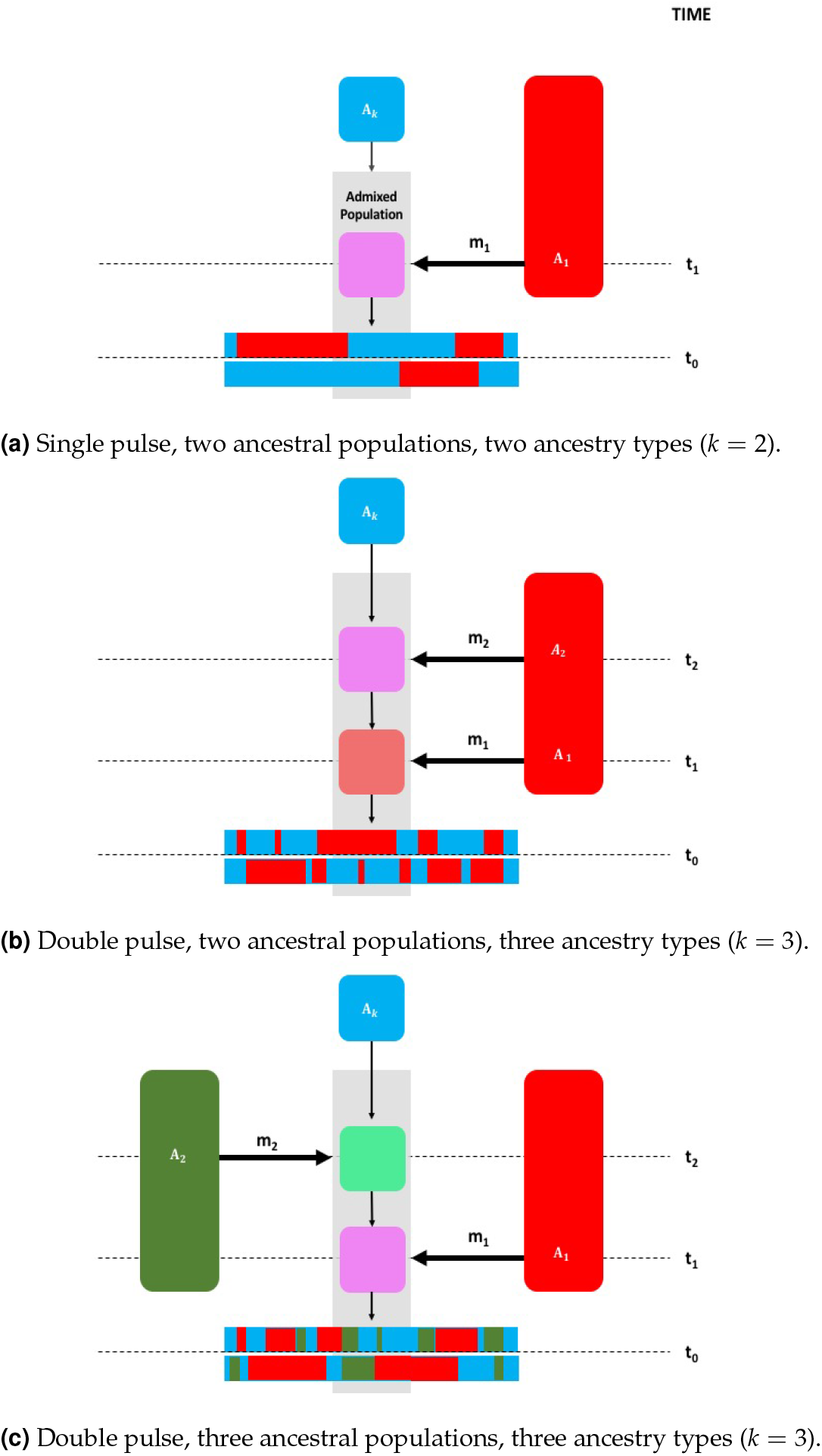
A schematic of (a) a single pulse model with two ancestry types, (b) a two pulse model with two ancestral populations, and (c) a two pulse model with three ancestral populations. *A_k_* is considered the ancestry type of the resident population and would be *A*_2_ in (a) and *A*_3_ in (b) and (c). Note that *A*_1_ and *A*_2_ may come from the same ancestral population, but are modeled as independent states. The gray shaded region draws attention to the admixed population(s). Time since admixture pulses are measured in generations and are denoted as *t*_1_ and *t*_2_, where *t*_1_ occurs more recently than *t*_2_. The time of sampling is represented by *t*_0_, where *t*_0_ = 0 if sampling occurred in the present. The proportion of ancestry in the admixed population that entered during an admixture pulse is denoted as *m*_1_ and *m*_2_. Colors represent genetically distinct ancestry types. Local ancestry across a chromosome after admixture is represented by horizontal bars at the bottom of each subplot.

Prior to this study, we developed a framework for simultaneously estimating admixture times and LA across the genomes of admixed populations (Corbett-Detig and Nielsen 2017). Our method can estimate LA and admixture times in a hidden Markov model (HMM) framework assuming a single admixture pulse model. Here, we extend this method to admixed populations that have experienced multiple ancestry pulses in their recent evolutionary past (Figure 1). We evaluate the performance of this approach using extensive forward-in-time admixture simulations. Finally, we apply this method to study admixed *Drosophila melanogaster* populations in sub-Saharan Africa, and we find evidence consistent with a recent single pulse of cosmopolitan admixture into African populations as well as evidence for more ancient admixture amongst genetically differentiated populations in sub-Saharan Africa.

## Model

### Overview

In our previous work, we developed an HMM approach to simultaneously estimate local ancestry and admixture times using next generation sequencing data in samples of arbitrary sample ploidy. No phasing is necessary, and unlike most LAI methods, ours models read pileups, rather than genotypes, making this approach appropriate for low coverage sequencing data (Skotte *et al*. 2013). Additionally, although not addressed in this work, our method accommodates samples of arbitrary ploidy, making it ideal for poolseq applications or for populations with unusual ploidy, e.g. tetraploids. Our previous work assumed a single exponential tract length distribution and is therefore limited to accommodating only a single admixture event between two populations (Corbett-Detig and Nielsen 2017). Here, we seek to extend this framework to accommodate additional admixture pulses either from distinct ancestry types or multiple pulses from the same type. Wherever possible, we have kept our notation identical to our previous work to facilitate comparisons between the models. Please refer to Table S1 for descriptions of the statistics we used to describe admixture model and LAI accuracy.

### Implementation and Availability

We implemented the following model into our software package called *Ancestry_HMM* (www.github.com/russcd/Ancestry_HMM). Below, wherever possible, we give the script name and line number responsible for a computation, denoted as *header: line_number*. All code is in the *src/* directory within the *Ancestry_HMM* repository. All line numbers and results reported here are based on version 0.94.

### State Space

Our model incorporates the ancestry of samples with arbitrary ploidy of *n* chromosomes and with *k* distinct ancestry types resulting from *k* − 1 admixture pulses. Therefore the state space *S*, is defined as the set of all possible *k*-tuples of non-negative integers, *H* = (*l*_1_,…,*l*_1_), such that 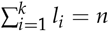 where *l_j_* is the number of chromosomes in the sample from admixture pulse *j*.

### Emission Probabilities

We model the probability of read pileup data (or alternatively genotypes) of each sample as a function of the allele counts in each ancestral population and assume a uniform prior on the allele frequencies in each source population. Note that where the same ancestral population contributes multiple pulses, the emission probabilities for each site at these states are identical and computed based on the sum of the number of chromosomes of each ancestry type. To accommodate up to *k* distinct ancestry types, we use multinomial read sampling probabilities. Specifically, if the representation in the read data is exactly equal (in expectation) for each chromosome, the probability of sampling a given read from chromosomes of ancestry pulse *k* is *l_k_/n*. Therefore, the probability of any given vector of read counts, *R* = (*r*_1_,…,*r_k_*), sampled from a site with depth *r* and across the chromosomes in a given hidden state *H* ∊ *S*, assuming no mapping or sequencing biases is (*read_emissions.h*: 31)

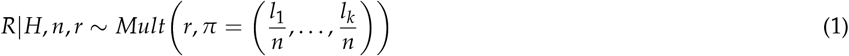

Conditional on the read count vector, *R*, the number of reads carrying the *A* allele (assuming an *A/a* di-allelic locus) is independent among ancestry pulses.

For each reference population, the allele count is binomially distributed given the (unknown) true allele frequencies, *f_j_*, *j* = 1 … *k*. Let *C_j_* represent the total allele count for reference population *j* and let *C_jA_* represent the total number of A alleles for reference population *j*. Then (*read_emissions.h*:51),

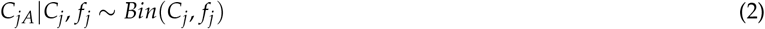

While we assume genotypic data for the reference populations, we assume short read pileup data for the admixed population. The (unobserved) allele counts in the admixed chromosomes, stratified by admixture pulse origin, *C*_*M*1*A*_, *C*_*M*2*A*_, … *C_MKA_* and *C*_*M*1*a*_, *C*_*M*2*a*_,… *C_Mka_* are also binomially distributed, i.e. for each ancestry type (*read_emissions.h*: 47),

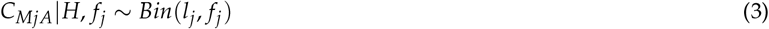

Then, assuming a symmetrical and identical error rate among alleles, *∊*, the probability of obtaining r_*jA*_ reads of allele *A* from within the r_*j*_ reads derived from chromosomes of type *j* is (*read_emissions.h*: 47),

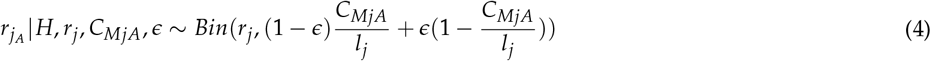

By integrating across all possible allele frequencies in the ancestral population assuming a uniform (0,1) prior, we obtain the probability of a given number of reads of allele *A*, r_*jA*_

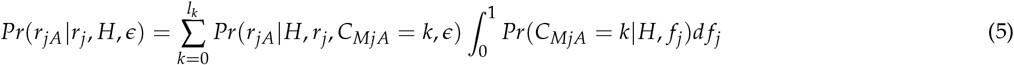

We then find all possible ways of arranging *r_A_* reads of allele *A* across the read vector *R, R_A_* = {(*r*_1*A*_,…,*r_kA_*) } Here, for each ancestry type, we require that 0 ≤ *r_jA_* ≤ *r_j_* and 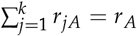. The probability of a given configuration is (*distribute_alleles.h*: 31),

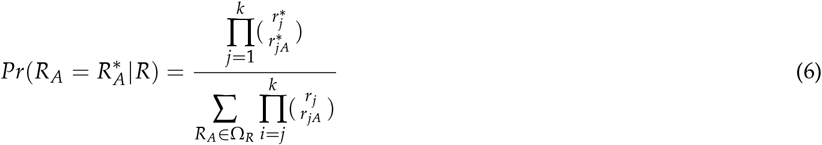

where Ω_*R*_ is the set of all configurations of *R_A_* compatible with *R*. For each configuration of reads, *R* and *R_A_*, these above probabilities for each ancestral population combine multiplicatively across all ancestral populations, and we are then able to obtain the emissions probabilities for the hidden state, *H* = *i*, as

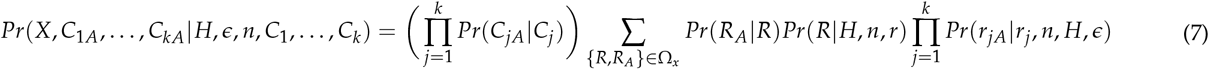

where *X* is all the read data for the admixed population for the site and Ω_*x*_ indicates the set of all values of {*R_A_, R*} compatible with *X*.

### Transition Probabilities

Transition probabilities must also be significantly expanded to accommodate the more complicated ancestry models investigated in this work.

### Modeling multiple pulses into a single recipient population

First, we consider a scenario where two ancestry pulses enter the same admixed populations at two distinct times (Figure 1c). Here, we will refer to the time of each ancestry pulse as *t_k_* where *t*_0_ = 0 and refers to the present and *t*_1_, for example, refers to the most recent admixture pulse in backwards time. During each pulse, a proportion of the resident population, *m_k_*, is replaced. In this model, there are two distinct epochs during which a recombination event may occur along a chromosome. The last epoch, in backwards time, occurs during the time interval between *t*_1_ and *t*_2_. During this time, there are two ancestry types present and the transition rate between them are identical as in our previous work (Corbett-Detig and Nielsen 2017). Specifically, the transition rate between the resident ancestry type *A*_3_ and ancestry type *A*_2_ is

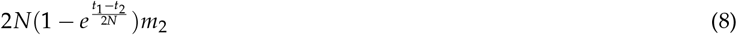

in units of Morgans per segment (Liang and Nielsen 2014). A nearly identical relationship holds for the transition rate from ancestry type *A*_2_ to *A*_3_,

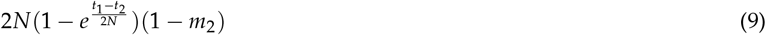

However, after the second ancestry pulse in forward time, additional transitions between ancestry types *A*_3_ and *A*_2_ will occur. During this interval, the transition rate from ancestry type *A*_3_ to ancestry type *A*_2_ is

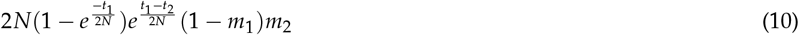

This transition rate reflects the chance of a recombination event occurring between *t*_1_ and the time of sampling with no back coalescence to the previous marginal genealogy in either time epoch. Finally, 1 — *m*_1_ is the probability that this recombination event does not choose a lineage that entered the population during the second ancestry pulse in forward time, and *m*_2_ is the probability of recombining with a lineage from the first ancestry pulse in forward time. Hence, the total rate of transition between ancestry type *A*_3_ and ancestry type *A*_2_ across both epochs, is

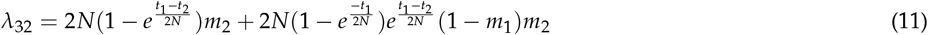

Again, a similar rate holds for transitions between ancestry types *A*_2_ and *A*_3_. Transition rates associated with the second ancestry pulse are simpler, and closely resemble those for a single pulse model. Specifically, the rate of transition from ancestry type *A*_3_ to ancestry type *A*_1_ is

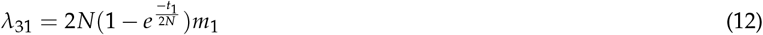

Here, we include the probability that a recombination event occurs within the second epoch and that the lineage selects ancestry type *A*_1_. Note that if the lineage selects ancestry type *A*_1_, we need not consider the probability of back coalescence in the time interval between *t*_1_ and *t*_2_ as it must exit the admixed population during the first ancestry pulse. Similarly, the transition rate from ancestry type *A*_1_ to ancestry type *A*_3_ is

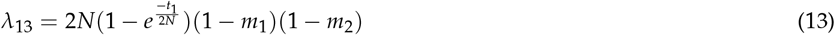

where this equation reflects the probability that a recombination event occurs during the second epoch and then the lineage selects ancestry type *A*_3_. Note that no consideration is given to back coalescence to the previous marginal genealogy during the epoch between *t*_2_ and *t*_1_ because there is no ancestry type *A*_1_ present in the population during this time. Here again, similar rates hold for transitions from ancestry type *A*_1_ to ancestry type *A*_2_ and from ancestry type *A*_2_ to ancestry type *A*_1_ following similar logic presented above.

Transition rates can be generalized to include an arbitrarily large number of distinct ancestry pulses. Briefly, transitions between ancestry types may occur during any epoch in which they are both present within the admixed population. For example, for the first pulse in forward time, transitions between ancestry type *A_k_* and *A*_*k*−1_ may occur anytime between *t*_*k*−1_ and the present. Therefore, all epochs will be necessarily included the transition calculations between ancestry type *A_k_* and *A*_*k*−1_. More generally, the transition rate between ancestry pulses, *i* and *j*, where 1 ≤ *i* < *j* ≤ *k* is,

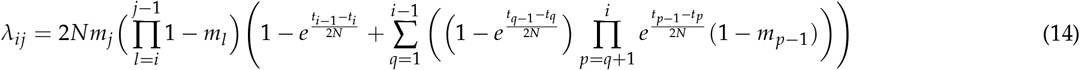

Conversely, if 1 ≤ *j* < *i* ≤ *k*, the transition rate between ancestry of pulse *i* to pulse *j* is very similar, i.e., transitions may occur during the same epochs (*t_j_* through the present). Therefore,

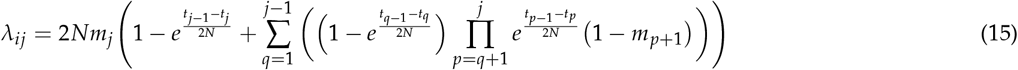

Note that it is not necessary for each ancestry pulse to be derived from distinct ancestral populations. Indeed, we conceived of this approach as a means of fitting multiple pulse ancestry models. Therefore any number of pulses may be contributed by as few as two ancestral populations. However, in order to model full transition rates, it is necessary to estimate the proportion contributed by each pulse even when they are from the same ancestry type, potentially making multiple pulse ancestry models more challenging to fit than models where each pulse is contributed by a separate subpopulation. More broadly, while this model is quite generalizable, there are limits to what is practical to infer using real datasets (see below).

### Transition Rates per Basepair

Equations 14 and 15 model transitions in Morgans per segment between ancestry states. We must therefore convert these expressions into a transition rate per basepair. To do this, we multiply the recombination rate by Morgans/bp using an estimate of the local recombination rate within that segment of the genome, *r_bp_*. Therefore, the single chromosome transition matrix, *P*(*l*) = *P_ij_*(*l*),*i,j* ∊ *S*, for a two pulse population model for two markers at distance l basepairs from one another would be as follows (*create_transition_rates.h* 69).

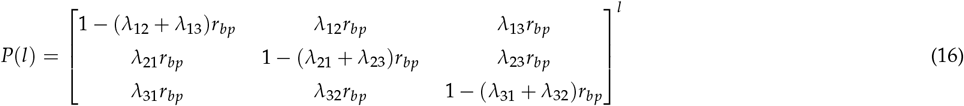

### Modeling Sample Ploidy

The above model describes the ancestry transitions along a single chromosome. However, many datasets contain samples that are diploid or pooled rather than haploid, or equivalently completely inbred. For simplicity, we model each sample of ploidy *n* as the union of n independent admixed chromosomes. To approximate the transition probability from state *i* to state *j* in a sample of *n* chromosomes, we assume the ancestry proportion, *m*, contributed by ancestry type *A* are known. Let the current hidden state ancestry vector be *s* = {*s*_1_, *s*_2_,…, *s_k_*} where *k* is the maximal number of ancestry types and *s_i_* indicates the number of chromosomes with ancestry component *i*. Then 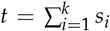 is the total number of different chromosomes in the pool. Furthermore, let *P_ij_*(*d*) be the *d*-step transition probability from ancestry *i* to *j* of the previously defined 1-chromosome process. Define *q_ij_* as *q_ij_* = *s_i_ P_ij_*(*d*). Also, let *a* = {*a*_1_, *a*_2_,…, *a_k_*} 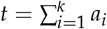, be the ancestry vector *d* sites downstream from the location of the locus with ancestry vector *s*. Then, we will approximate the transition probability from, *s* to *a*, as

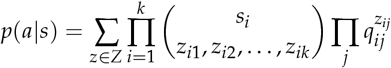

where *z* = {*z_ij_*} is an *k* × *k* matrix of non-negative integers and *Z* is the set of all such matrices for which 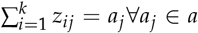 and 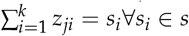. The sum over all *z* ∊ *Z* can be large and increases exponentially in *k*.

The true ancestral recombination graph is potentially more complex than our simple approximation which assumes all ancestry transitions to be independent. Therefore, caution is warranted when using our model on samples with higher ploidy (Corbett-Detig and Nielsen 2017) (*transition_information.h*: 21).

### Model Optimization

Because the search space for admixture models is potentially quite complex, we have implemented optimization using the Nelder-Mead direct search simplex algorithm (Nelder and Mead 1965) to optimize the HMM using the forward-equation to compute model likelihoods. As with all direct search algorithms, there is no guarantee that the optimum discovered is a globally optimal solution. We therefore include random-restarts to insure that the globally optimal solution can be recovered consistently.

### Assumptions

Perhaps the central assumption of this approach is that admixture occurs in discrete, distinct “pulses”. Whereas this is violated in a wide array of true admixture events with either ongoing or periodic admixture (Pool and Nielsen 2009; Gravel 2012), the pulse model is tractable for estimating admixture histories (Corbett-Detig and Nielsen 2017; Loh *et al*. 2013; Gravel 2012). We have previously shown that LAI using our method is robust to a wide array of perturbations including continuous migration and natural selection (Corbett-Detig and Nielsen 2017). Nonetheless, all results, particularly those that hinge heavily on the precise timing of admixture events, should be interpreted cautiously. See also our discussion below.

## Validation

### Confirmation of the Ancestry Tract Length Approximation

We first confirmed that our sequential Markov coalescent (SMC’) approximation for the ancestry tract length distribution is correct. Specifically, we simulated tract length distributions from the forward-in-time simulation program, SELAM (Corbett-Detig and Jones 2016), and compared those with the expected tract length distribution under our model. In comparing the two, we found that the model provides an excellent approximation for the ancestry tract length distribution (Figure 2). This therefore indicates that our SMC’ tract length approximation is likely to be sufficient for our purposes. Moreover, we also confirmed that this framework can also accurately accommodate models involving more ancestral populations (Figure S1, Table S2).

### Simulation Ancestral Populations

We used the approach of Corbett-Detig and Nielsen (2017) to validate our HMM software implementation and to test the performance of this expanded model. Briefly, we simulated ancestral genotype data using the coalescent simulation framework MACS (Chen *et al*. 2009), we then simulated ancestry tract length distributions in forward-time using our software package SELAM (Corbett-Detig and Jones 2016). Genotype data for admixed individuals was then drawn from the ancestral data simulated using MACS where genotypes were drawn from each population and assigned to each tract based on that tracts’ ancestry type. To explore a wide range of genetic divergences among ancestral populations, we simulated genotype data for ancestral populations at varying levels of genetic divergence from one another. Specifically, we considered populations that are 0.05, 0.1, 0.25, 0.5, and 1 N_*e*_ generations divergent from one another.

### Linkage Disequilibrium (LD) Pruning

LD among sites may inflate transition rates between ancestry states. In order to mitigate this, we first pruned sites in strong linkage disequilibrium in each reference panel by computing LD among all pairs of markers within 0.1cM of each other. We then discarded one site from each pair with increasingly stringent pruning until we found that admixture time estimates were approximately unbiased in two pulse admixture models. We note that the LD pruning necessary for two pulse models appears to be sufficient for fitting more complex models of admixture (see below). This suggests that simple single-pulse admixture simulations could be used successfully to determine the necessary levels of LD pruning. Also, we found that substantially more stringent LD pruning is necessary to produce unbiased admixture time estimates for ancestral populations that are minimally divergent from one another (Figure S2).

### Three Ancestral Populations

We next sought to evaluate the accuracy of this method when two ancestry pulses are contributed from distinct ancestral populations. For all levels of population divergence considered, we find that our method recovers the correct times of admixture events with reasonably high accuracy for recent admixture times (Figure 3). Notably, the error in time estimates decreases substantially with increasingly divergent ancestral populations indicating that this method will produce the most accurate results when ancestral populations are highly genetically divergent.

**Figure 2.**
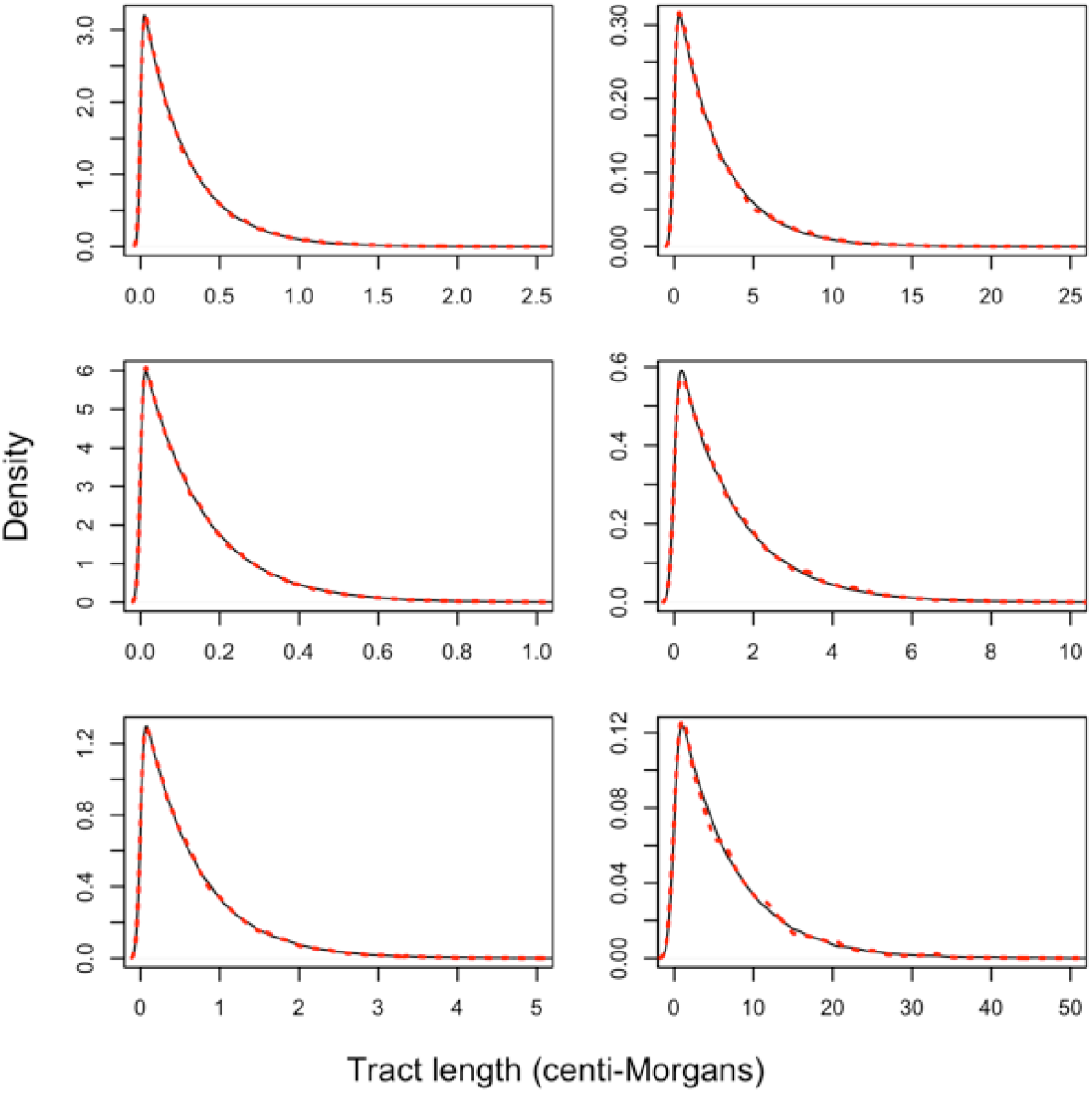
Tract length distributions obtained using our tract length model approximation (solid black) and forward-in-time simulation (dashed red). A model like Figure 1c was considered. An initial pulse of ancestry type *A*_2_ entered a resident population of ancestry type *A*_3_ and replaced 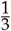 of the resident population. Then, a second ancestry pulse in forward time, from ancestry type *A*_1_, replaced 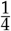 of the resident population. Each simulation had a diploid population size of size 10,000 and we aggregated data from 50 sampled individuals across 100 simulations to produce the full tract length distribution. From top to bottom, respective ancestry tract length distributions correspond to ancestry type *A*_3_, *A*_1_, and *A*_2_. We investigated two admixture models. The first model has a first and second pulse occurring 200 and 1,000 generations, respectively, before the present (left). The second model has a first and second pulse occurring 20 and 100 generations, respectively, prior to the present (right). Note that axes differ between figure panels.

Whereas our method accurately estimates admixture times when admixture events occur at intermediate times prior to sampling, we find that the accuracy suffers somewhat when estimating the timings of two more ancient admixture events. Indeed, for modestly divergent ancestral populations, e.g., 0.1N_*e*_ generations divergent, we find that our approach consistently underestimates the times since admixture events, particularly for the more ancient ancestry pulse (Figure 3). This is true despite a nearly linear relationship for estimated admixture times and actual admixture times in a single pulse two ancestral population model (Figure S2), indicating that there is a significant cost for accuracy of model estimation when estimating additional ancestry pulse parameters. However, for more divergent ancestral populations, it is still feasible to obtain accurate admixture time estimates (Figure 3). We note that our good performance in estimating old admixture times may be a function of more informative sites and less LD present in *Drosophilα melanogaster* compared to humans, for example. Additionally, because there is still a monotonic relationship between the time estimates and true admixture times, it might be feasible to correct for this bias and obtain accurate admixture time estimates even when ancestral populations are not particularly divergent.

### Two Pulse Admixture Model

It is significantly more challenging to distinguish between models with a single ancestry pulse and models with two distinct ancestry pulses from the same ancestral population. As we described above, one of the key difficulties is that in addition to estimating admixture times, our model must also estimate the proportion of the total ancestry contributed by each pulse. Furthermore, it is essential that this approach includes a mechanism for distinguishing between single and double pulse models. We therefore began with a single-pulse admixture simulation and fit both single and two pulse admixture models to these data.

We find that traditional likelihood ratio tests (LRT) universally favor two pulse versus single pulse models even when data are simulated under a single pulse model for all scenarios that we considered (Figure S3). Additionally, we note that two-pulse models may become degenerate, either because a pulse’s ancestry proportion may be nearly zero or because two pulses may occur at virtually identical times. We therefore caution that these factors could make traditional LRTs challenging to interpret. We therefore suggest that when using this method to analyze admixed populations, it will usually be preferable to choose the simplest admixture model that is consistent with the data. More complex models are usually favorable by standard statistical comparisons, but may overfit the data. This consideration is reminiscent of concerns for the STRUCTURE model (Pritchard *et al*. 2000; Evanno *et al*. 2005). In general, this means the model that contains the fewest distinct ancestry pulses should be selected.

**Figure 3.**
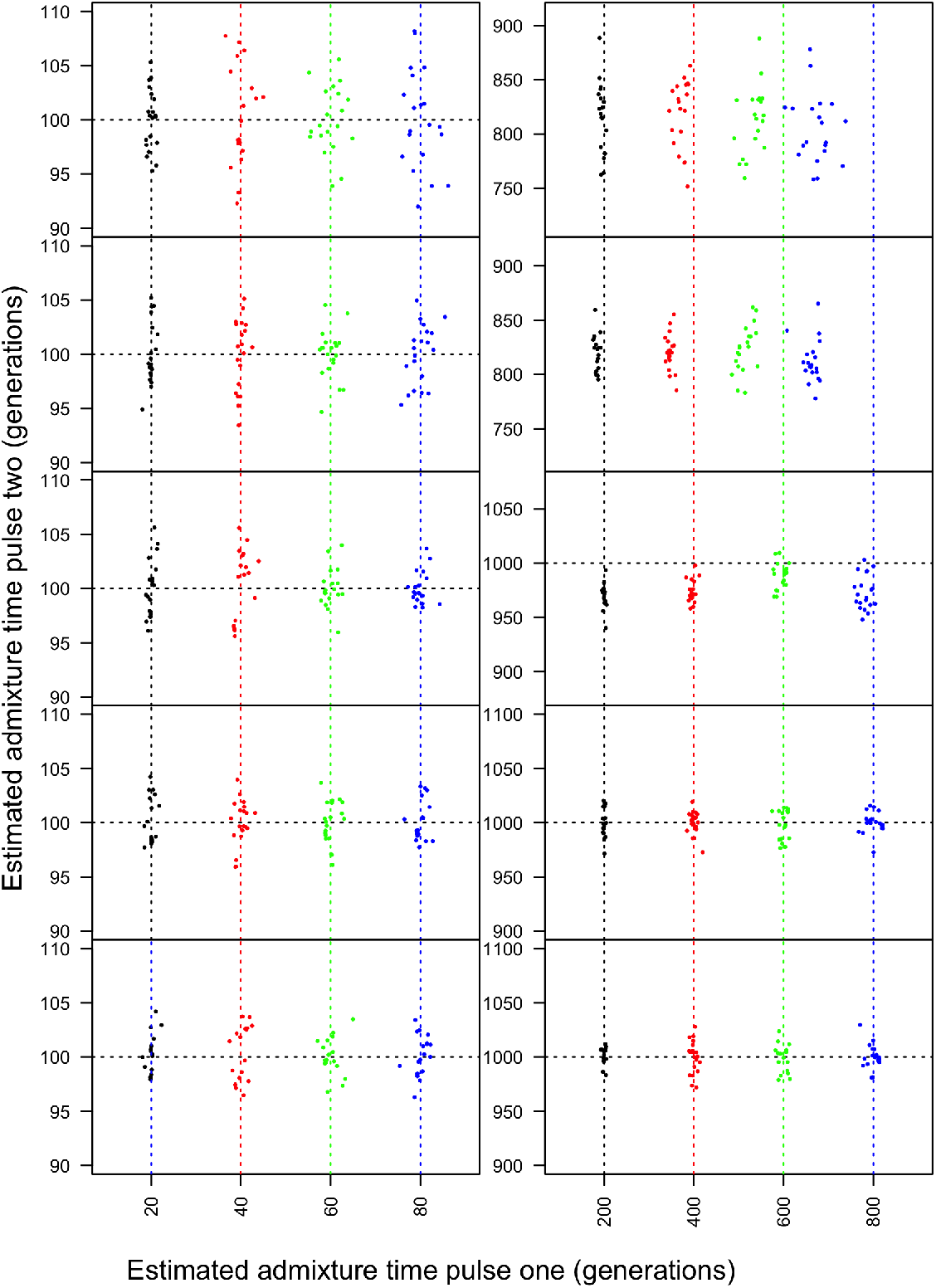
Admixture time estimates for two pulse population models with three distinct ancestral populations. From top to bottom, panels include the divergence time between the ancestral populations is 0.05, 0.1, 0.25, 0.5 and 1 N_*e*_ generations. True admixture times are indicated by the dashed lines. On the left, for all admixture models considered, the second pulse occurred 100 generations before the present. The first pulse occurred 20 (black), 40 (red), 60 (green), and 80 (blue) generations prior to sampling. On the right, the second pulse occurred 1000 generations before the present and the first pulse occurred at 200 (black), 400 (red), 600 (green) and 800 (blue) generations prior to sampling. Note that due to underestimation of admixture times, the figure axes differ between panels for plots of more ancient admixture events.

Spurious admixture models can be identified by interrogating the timing and proportion of ancestry contributed by each admixture pulse. First, many of the resulting two pulse admixture models contain ancestry pulses with similar times (Figure 4, Tables S3, S4, S5), suggesting that these could be recognized as representing a single admixture event. Second, of the models containing two pulses, many contain one pulse near the correct admixture time and another distant pulse that introduced a very small proportion, i.e. 1% or less, of the total ancestry in the sampled admixed population (Figure 4, Tables S3, S4, S5). Often the pulse introducing a small proportion of ancestry in the sampled population occurred in the distant past, where admixture time is harder to estimate. Therefore, admixture pulses that (1) occur in the distant past and contribute relatively small proportions of the total ancestry or (2) pulse at similar times, may indicate a spurious admixture model and should be disregarded in favor of simpler admixture scenarios.

When we simulated admixed populations whose histories contained two distinct ancestry pulses, we found two-pulse models could sometimes be fit accurately. Specifically, when the two ancestry pulses occurred at fairly different times (e.g., 20 and 100 and 200 and 1000 generations), our approach identified models that correspond closely to the parameters under which the admixed population was simulated. However, when the two ancestry pulses occurred at closer time intervals, we were less frequently able to reliably recover the correct admixture model (Figure 5, Table S4). Additionally, we note that the accuracy of two-pulse models depends on the genetic distance between ancestral populations (Figure 5), where it is generally more straightforward to fit admixture models for genetically distant ancestral populations.

**Figure 4.**
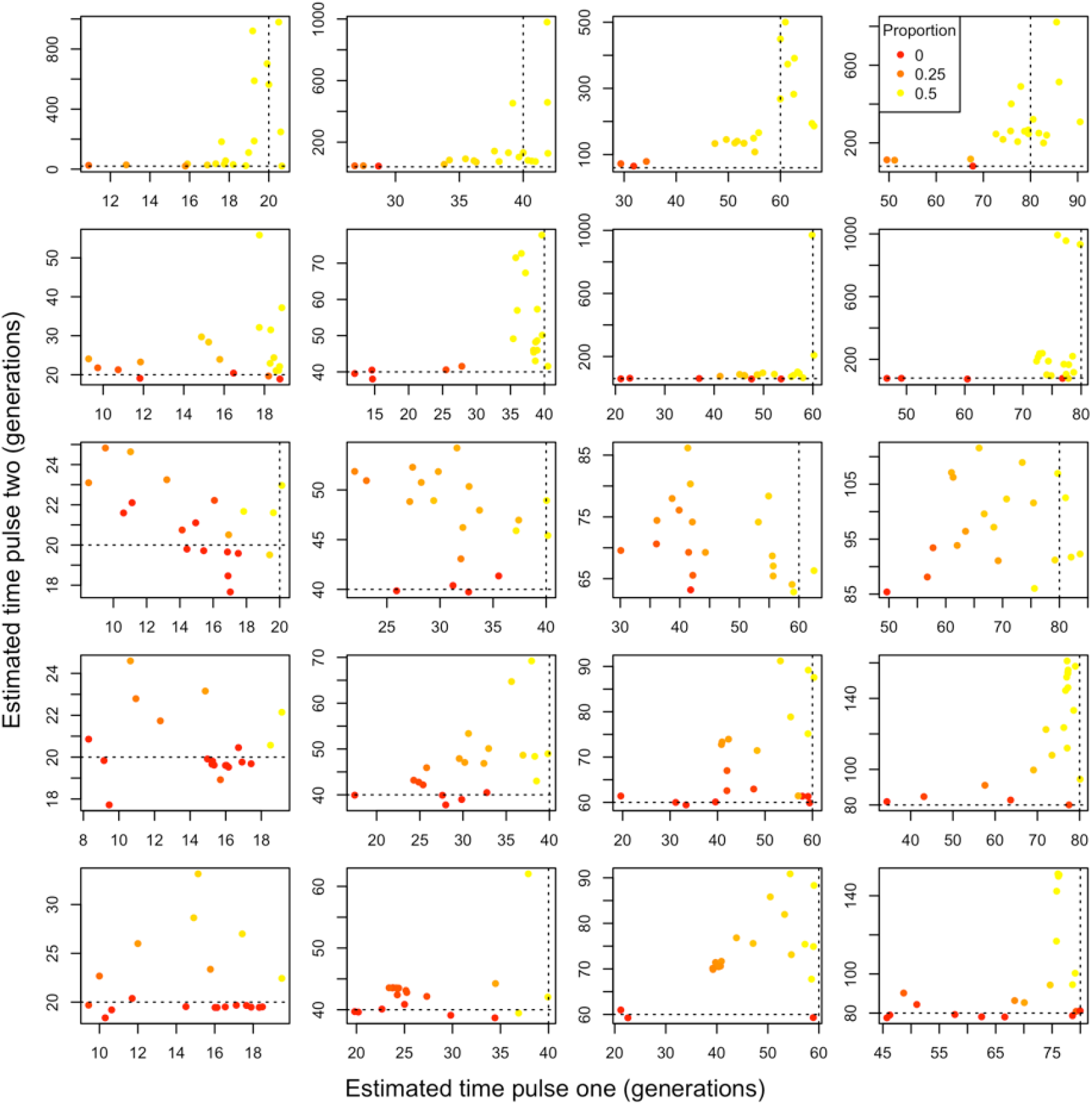
Two-pulse admixture models fitted using our framework to data generated under a single pulse admixture model. We considered varying levels of population divergence. From top to bottom, the ancestral populations are 0.05, 0.1, 0.25, 0.5, and 1 N_*e*_ generation divergent from one another. From left to right, the single admixture pulse occurred 20, 40, 60, and 80 generations prior to sampling and replaced one half of the ancestry. Point colors correspond to the proportion of ancestry that is attributable to the second pulse with a gradient running from 0 (red) to the maximum possible, 0.5 (yellow). Dashed lines reflect the true admixture time. Note that axes differ between figure panels.

Collectively, our results suggest that it might not be possible to distinguish between single-pulse and two-pulse admixture models when ancestry pulses occurred at similar times. However, it is feasible to distinguish single pulse models from admixture models with both relatively ancient and recent admixture. Therefore, we anticipate that this approach will be valuable for investigating a range of hypotheses with dramatically different admixture times. Furthermore, as we consider here intermediate-sized population samples, i.e. 50 individuals, it may be feasible to distinguish more fine-grained admixture models using larger samples sizes.

### LAI Accuracy

We note that while two pulse models do improve the accuracy of LAI, the improvement observed tends to be slight (Tables S3, S4, S5). This suggests that for studies aimed at evaluating patterns of LA across the genome, there may be no need to extensively optimize admixture models to obtain reasonable estimates of the LA landscape in the admixed population. Single pulse models may be sufficient to accurately characterize LA across the genome for many admixed populations.

### Robustness of our Approach to Smaller Sample Sizes, Divergent Reference Populations, and Inaccuracies in the Genetic Map

We anticipate our approach will be used to estimate the timing of admixture in model and non-model systems. It is likely that the underlying assumptions will be violated in some applications. To characterize our model’s performance and robustness in light of these challenges, we evaluated our method’s ability to correctly estimate the timing of admixture under varying sample sizes, reference population divergences, and erroneous recombination maps. See S1 for definitions of summary statistics used to quantify accuracy of admixture models and LAI.

**Figure 5.**
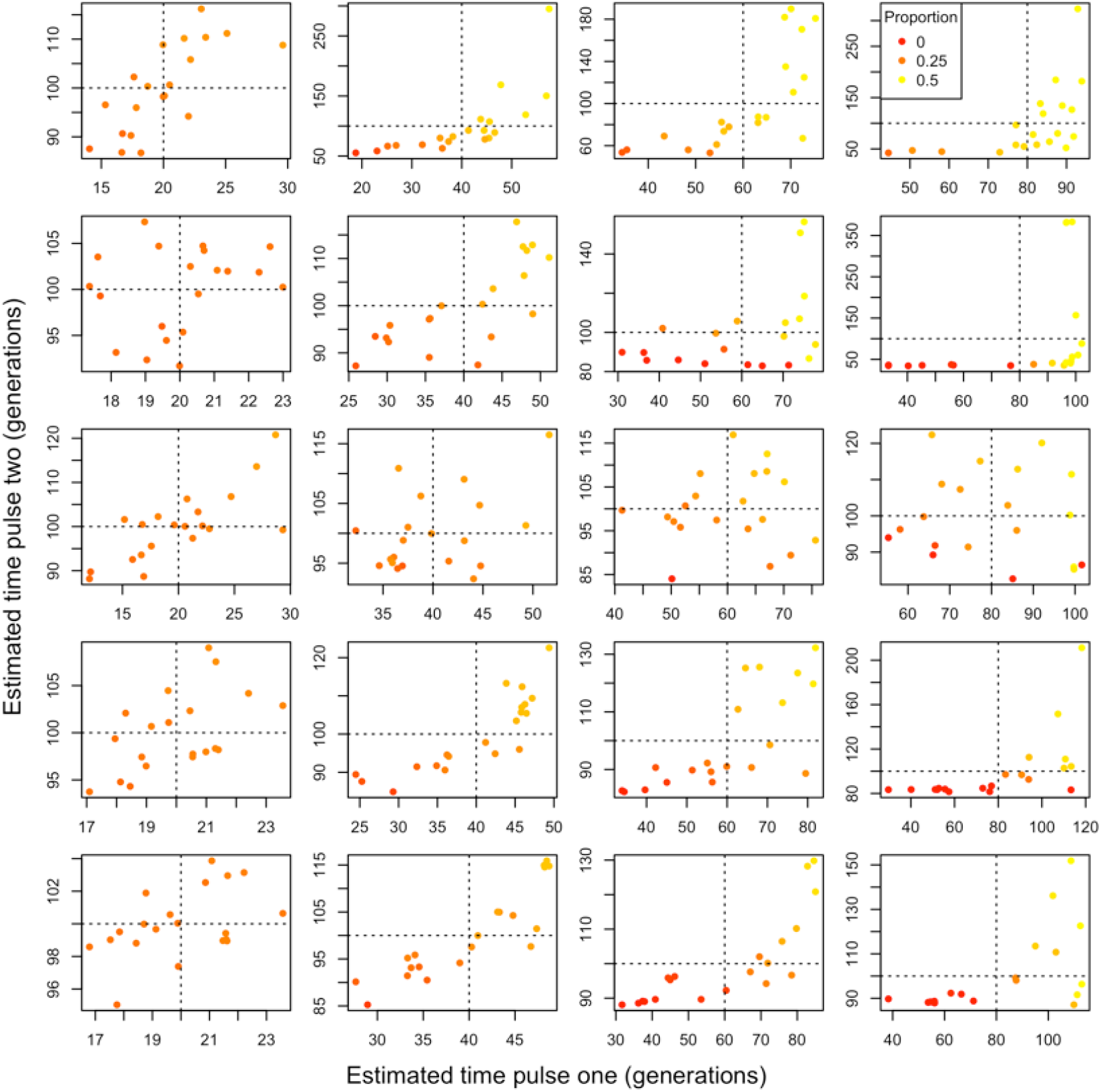
Two-pulse admixture models fitted using our framework to data generated under a two pulse admixture model. We considered varying levels of population divergence. From top to bottom, the ancestral populations are 0.05, 0.1, 0.25, 0.5, and 1 N_*e*_ generation divergent from one another. For all models, the second admixture pulse occurred 100 generations prior to sampling. From left to right, the first pulse occurred 20, 40, 60, and 80 generations prior to sampling. The second pulse replaced 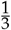 of the resident population and the first pulse replaced 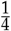 of the resident population. Therefore, each ancestral population contributed one half of the ancestry at the time of sampling. Point colors correspond to the proportion of total ancestry at the time of sampling that is attributable to the first pulse with a gradient running from 0 (red) to the maximum, 0.5 (yellow). Dashed lines reflect the true admixture times for the first and second pulse. Note that axes may differ between subplots.

Accurate fine-scale recombination maps are necessary for estimating transition rates between adjacent ancestry states in our model. However, we anticipate that our method will be applied to non-model systems where a fine-scale recombination map might not be available. Therefore, We tested the resilience of our method against perturbations of the recombination map using populations that are 0.25 Ne generations divergent (Methods). The normalized root mean squared error (NRMSE) in estimating the timing of a single pulse admixture event using our approach is less than 12% for all admixture times 20, 40, 60, and 80 even when the recombination map is substantially distorted (Figure S5, Tables S6 and S7). Moreover, our approach is unlikely to give false support for a two pulse model to truly single pulse data (Figure S6). When fitting a two pulse model to two pulse data, our method consistently over-estimates the timing of admixture events when the true recombination map is distorted relative to the assumed map. However, for all but the most strongly distorted recombination maps our results imply that two pulse models can be reliably identified, albeit with overestimated admixture times (Figure S7). Therefore, for single pulse admixture models, a high density recombination map does not appear necessary, however, for accurate two pulse admixture models, a high density genetic map may be required to obtain accurate admixture time estimates. See Table S7 for detailed error reports as a result of perturbed rates of recombination.

Additionally, the source population contributing to the admixture history of a population may not always be available for genetic sequencing, or may be incorrectly identified. To test the sensitivity of our approach when a related but distinct population from the true admixture source is used as a reference, we simulated admixture events using reference populations of varying divergence to the actual source population and estimated the timing of admixture (See Methods). We found our approach consistently overestimated the timing of admixture in both single and double pulse models (Figures S8, S9). In general as reference populations are more divergent from the true admixture source, our method increasingly overestimated the timing of admixture events. Nonetheless, it is feasible to identify a two-pulse model, despite overestimating the timing of admixture (Figure S9). Moreover, we fitted a double pulse model to single pulse data and found no false support for a double pulse model (Figure S10). We advise that when using our approach it will be necessary to carefully identify the closest possible reference population, as there are limitations of how divergent the reference can be to the actual source population in order to obtain accurate estimates of admixture time (Table S8, S9).

Lastly, we evaluated our method’s performance under varying sample sizes of the admixed population. Because we considered only a single chromosome in all analysis, we note that our sample sizes considered here, down to 10 individuals, are very modest relative to most sequencing applications with respect to total data considered. Regardless, we do not obtain false support for a two pulse model when applied to admixed populations that experienced only a single pulse (Figure S11). Estimating the timing of admixture becomes more accurate with large sample sizes. However, a sample size of just 10 individuals at a single chromosome still has a NRMSE less than 6% for all admixture times we considered (Tables S10, S11, S12). Therefore, we conclude that our approach is still useful in cases with limited sampling.

### Comparison to ALDER

We compared the accuracy of our approach in estimating the timing of admixture to that of ALDER (Loh *et al*. 2013). Briefly, ALDER and its extension, MALDER (implemented in Pickrell *et al*. (2014)), is based on modeling the decay of admixture LD within admixed populations. In all single and double pulse models that we considered (See Methods), our approach more accurately estimates the timing of admixture (Tables S13, S14). Notably, our method significantly outperforms ALDER in models involving pulses occurring more distantly in the past (i.e. more than 80 generations ago). This discrepancy might be attributable to our method’s ability to model short tracts of LA, which get disregarded in ALDER by default (Loh *et al*. 2013), potentially impacting the ability to identify ancient admixture events (Figure S12). Considering a relatively recent two pulse model where *t*_1_ = 20 and *t*_2_ = 100, MALDER correctly fits a two pulse model 6 out of 20 of the simulations, but selects a single pulse model for the 14 others (Table S14). Our approach fit a two pulse model to the model in all 20 simulations and produced a NRMSE of 0.0307 and 0.0214 for *t*_1_ and *t*_2_, respectively, compared to MALDER’s 0.1207 and 0.1856. Consistent with our results above, neither MALDER nor our method was able to consistently fit two pulse models when admixture events occurred more closely together in time (Tables S13, S14).

### When Is This Method a Good Choice?

Our software performs best with reference populations that are minimally divergent to the source population and an accurate fine-scale recombination map. Our results suggest that this approach is robust to varying levels of data availability and to errors in the fine-scale genetic map. Given that this is the only approach for LAI and admixture time estimation that can accommodate non-diploid samples, short read pileup data, and relatively ancient admixture pulses, it will likely be useful in a variety of systems.

Nonetheless, we advise carefully considering, *e.g*., through simulation, how this approach will perform with available data particularly when ancestral LD is extensive or populations are only weakly genetically differentiated, such as what is found in some human admixed human populations. Particularly for these situations, we suggest to instead apply LAI software that explicitly models the haplotype structure of admixed samples (Maples *et al*. 2013; Price *et al*. 2009; Dias-Alves *et al*. 2018) and use the resulting tract length distribution to estimate admixture time (Gravel 2012; Ni *et al*. 2018).

## Application

### Admixture in D. melanogaster

Biogeographical evidence (Lachaise *et al*. 1988) and patterns of genetic variation (Thornton and Andolfatto 2006; Begun and Aquadro 1993; Caracristi and Schlötterer 2003) suggest that *D. melanogaster* originated in sub-Saharan Africa, and went on to colonize much of the rest of the world relatively recently (Duchen *et al*. 2013). During this expansion, the population that left sub-Saharan Africa experienced a dramatic bottleneck that reshaped patterns of genetic diversity across much of the genome (Caracristi and Schlötterer 2003; Ometto *et al*. 2005; Duchen *et al*. 2013). The resulting “cosmopolitan” haplotypes are distinguishable from those of the ancestral sub-Saharan populations based solely on patterns of genetic variation (Pool *et al*. 2012) and these two ancestral populations have since encountered each other and admixed in numerous geographic locations within sub-Saharan Africa and worldwide (Pool *et al*. 2012; Schlötterer 2003).

Thus, *D. melanogaster* has emerged as an important model for understanding the genomic and phenotypic consequences of admixture in natural populations. There are important pre-mating behavioral isolation barriers between cosmopolitan and African individuals (Ting *et al*. 2001), as well as substantial phenotypic consequences that operate post mating within admixed individuals (Lachance and True 2010; Kao *et al*. 2015). Additionally, recent work has investigated the genomic consequences of admixture both demographically (Duchen *et al*. 2013; Bergland *et al*. 2016) and with the goal of evaluating the impact of natural selection in shaping genome-wide patterns of variation (Pool 2015). The genomic consequences of admixture have only been studied using relatively simple demographic models (i.e. a single event, (Pool *et al*. 2012)), leaving open the possibility that more complex admixture dynamics are common in natural populations of this species. Accurately characterizing the admixture histories and the patterns of local ancestry across the genome is essential to further our understanding of the demographic history of this species and build a complete null model for studying natural selection on ancestry during admixture.

### Evaluating Possible Applications to D. melanogaster Admixed Populations

In agreement with our general conclusions from admixture simulations, we find that our method is accurate, and single-pulse admixture models are distinguishable from two-pulse models for temporally distinct admixture events (i.e., wherein two pulses occurred at significantly different times, Figure 6). However, also consistent with our results above, we find that when admixture pulses occurred at relatively similar times (e.g. *t*_1_ = 200 and *t*_2_ = 250 generations prior to sampling), a single ancestry pulse contributes the vast majority of admixed ancestry, and this scenario is therefore ultimately most similar to a single pulse admixture model. These data therefore suggest that it is possible to distinguish single from two pulse admixture models in data consistent with *D. melanogaster* ancestral populations when the admixture pulses occurred at dramatically different times.

Perhaps owing to the higher marker density, the greater accuracy of genotyped markers along the genome, and/or the inbred genomes in our Drosophila samples, we find that our method is slightly more accurate for population histories consistent with those of *D. melanogaster* than for the simulated populations considered above (Tables S3, S4, S5). Additionally, admixture times can be accurately estimated even when admixture events occurred in the distant past (e.g., 5000 generations prior to sampling, Figure 6).

**Figure 6.**
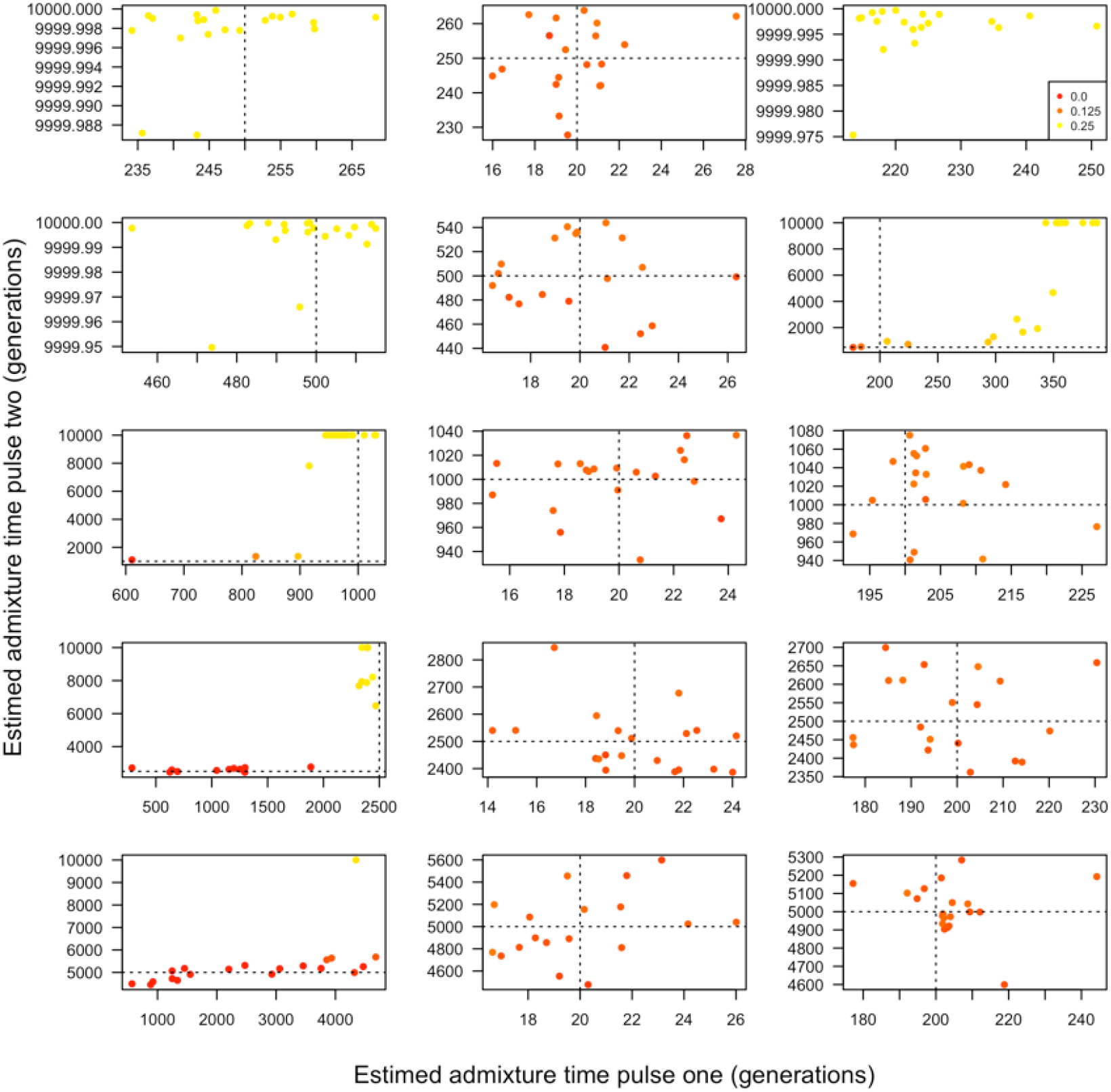
Admixture model fitted for data consistent with admixed Drosophila populations. Two-pulse admixture models for scenarios that are truly single-pulse (left), two pulse with a first admixture pulse 20 generations prior to sample (middle), and two pulse models with a first admixture pulse 200 generations prior to sampling (right). From top to bottom, the second admixture pulse occurred 250, 500,1000, 2500, and 5000 generations prior to sampling. The point colors indicate the proportion of ancestry in the sampled population that entered during the first admixture pulse in backwards time with a gradient running from 0 (red) to the maximum, 0.5 (yellow). Dashed lines indicate the correct timing of simulated ancestry pulses. In two pulse models, the first pulse contained 10% of the final ancestry and the second contributed 14% of ancestry to the sampled population. In single pulse models, all of the non-native ancestry is contributed by one pulse. These values were selected to be consistent with those from admixed sub-Saharan populations of this species (Pool *et al*. 2012). Note that axes may differ between subplots.

### Application to Admixed D. melanogaster Samples

To demonstrate the performance of our method on real datasets, we applied our approach to *D. melanogaster* variation data from sub-Saharan African populations (Figure 7). We studied populations from Rwanda (RG), South Africa (SD) and Gambella, Ethiopia (EA) and estimated the time of cosmopolitan admixture using both double and single pulse models.

In two pulse models, cosmopolitan ancestry pulsed twice (Figure 1b) and resulted in the most recent pulse contributing approximately 99% and 97.5% (for RG and SD respectively) of the total cosmopolitan ancestry present in the population at the time of sampling. The second, more ancient pulse, tends towards the maximum time allowed in our inference method, 10,000 generations, and contributes 0.01% and 2.5% of the total cosmopolitan ancestry present in the population at the time of sampling RG and SD (Figure 7). As described above, a distant admixture pulse contributing small amounts of ancestry likely indicates a spurious admixture model. Though it is possible that ancient cosmopolitan admixture contributed a small amount to the ancestry of the SD and RG populations, the simplest model, a single pulse model with cosmopolitan ancestry pulsing into African ancestry, provides a reasonable description of the demographic history of RG and SD. The estimated admixture times, 140 and 437 generations in a single pulse model, suggest that the observed cosmopolitan ancestry has entered these populations exceptionally recently despite earlier contact and extensive commerce among human groups.

To test how our method performs when ancestral populations are omitted, we applied a two pulse model to genotype data from Gambella, Ethiopia (EA) which is believed to have West African, Ethiopian, and cosmopolitan ancestry (Lack *et al*. 2015). Instead of modeling all three ancestral populations, we omitted West African as an ancestral population to mimic a scenario where the ancestral populations of an admixed population have not all been identified. The results of the two ancestral state model estimated an ancient admixture pulse composing 6.5% of the total cosmopolitan ancestry present in the population at the time of sampling (Figure 7). Importantly, we note that 6.5% is substantially more than we found in any of our simulated single-pulse datasets. A second pulse about 200 generations ago contributed 28% of cosmopolitan ancestry. Since this two pulse model did not include ancestry from a West African population, it is likely that our method fit a portion of the allele frequency differentiated between the ancestral populations into the ancient pulse of cosmopolitan ancestry. Thus, a large proportion of ancestry contributed by an ancient admixture pulse might indicate the need to identify additional ancestral populations. A two pulse model including three ancestral populations is therefore more appropriate for studying the admixture history of EA.

**Figure 7.**
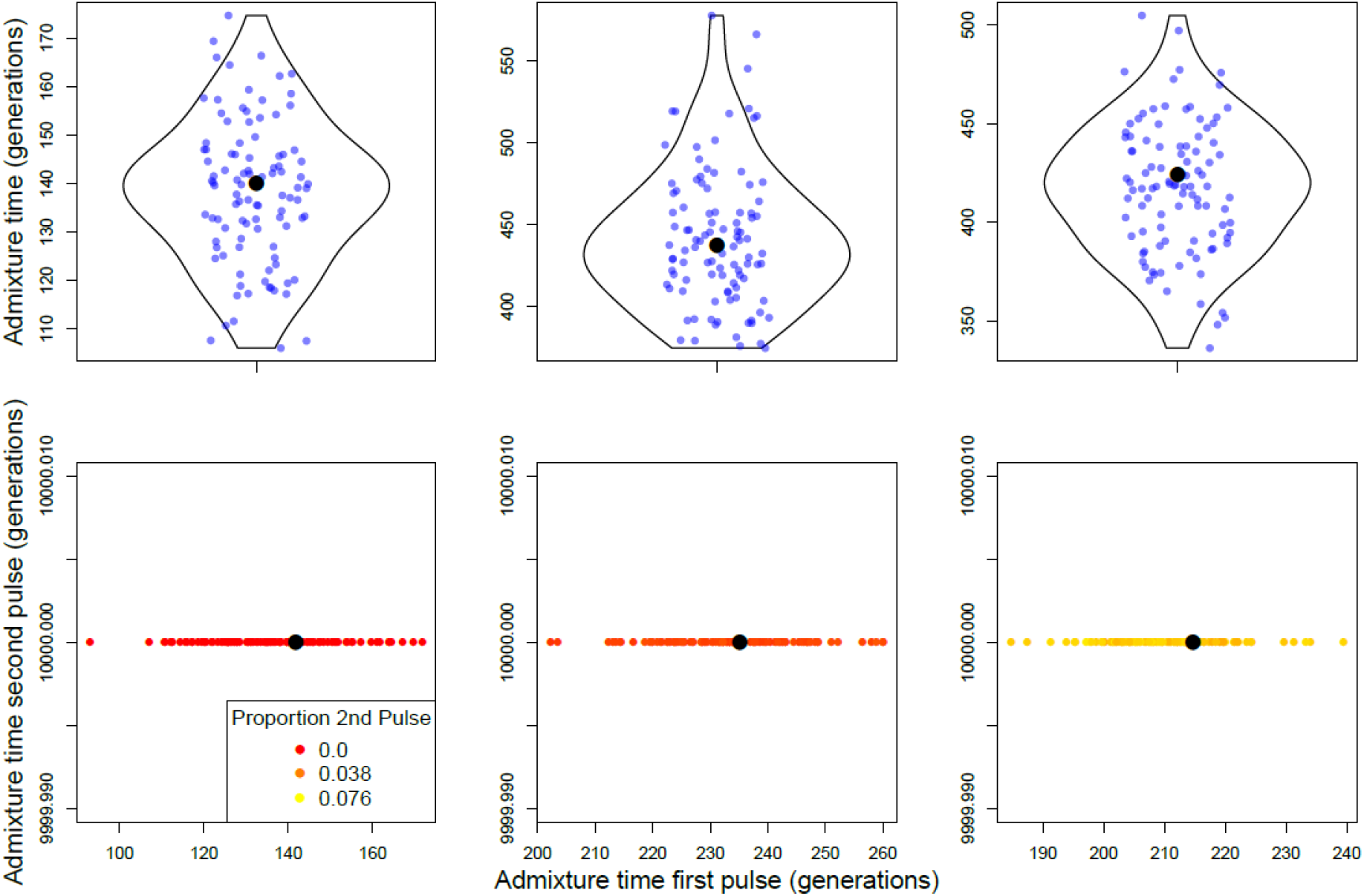
Single and double pulse models applied to real genotype data generated from sub-Saharan African populations of *Drosophila melanogaster*. In the plots shown, cosmopolitan ancestry pulsed into native African ancestry once (top row) or twice (bottom row). The panels from left to right are populations RG, SD, and EA. Single and double pulse models were bootstrapped 100 times. Each point in the top row corresponds to a bootstrap estimate of the timing of admixture. The bottom row shows bootstrap estimates for the timing of the second admixture pulse in backwards time. The point colors indicate the proportion of ancestry in the sampled population that entered during the second (in backwards time) admixture pulse with a gradient running from 0 (red) to the maximum, 0.076 (yellow). Black dots indicate the bootstrapped time optima. Note that axes may differ between subplots.

### Application to Three Population Mixture in Gambella, Ethiopia (EA)

Our results above emphasize the importance of identifying and incorporating variation data from all ancestral populations. Here, we use a two pulse, three ancestral population, admixture model to estimate the order and timing of admixture events into EA (Figure 1c, Figure 8).

The native ancestry of *Drosophila melanogaster* from EA is unknown. Previous studies have suggested both cosmopolitan and sub-Saharan ancestry contribute to the genetic variation in EA (Figure 1c) (Lack *et al*. 2015). To determine the most likely native ancestry of EA, we applied three permutations of a two pulse model to population data from Gambella, Ethiopia. We used unadmixed populations from Ethiopia (EF), cosmopolitan (FR), and West African (AF) to represent ancestral populations of EA and used our program to estimate the timing of admixture in each admixture model. Our three models were: EF and AF pulsing into resident FR ancestry, AF and FR pulsing into EF ancestry, and EF and FR pulsing into AF ancestry.

When FR is supplied as the resident population in the two pulse model, EF and AF both enter the population 1140 generations ago. This is implausible for a variety of reasons. First, even without considering this result, it is unlikely that cosmopolitan ancestry is the native ancestry type in Gambella, Ethiopia. Second, it is exceptionally unlikely that two distinct ancestral populations enter the admixed population at precisely the same time. Finally, the likelihood of the FR native ancestry model is less favored relative to an EF or AF native model by 11000 log likelihood units. We therefore consider a model with native African ancestry, EF or AF, to most likely describe the admixture history of Gambella, Ethiopia *D. melanogaster*.

When EF and AF were modeled as native ancestry, resulting admixture histories were nearly identical, suggesting either EF or AF could be native to Gambella since these are equivalent models. Using a two pulse, three ancestral population model, we estimated the timing of EF and FR admixture pulses in AF native ancestry (Figure 1c). The second pulse in backwards time is estimated to have occurred approximately 4962, 95% CI [4349, 5619], generations ago and suggests that EA admixed with genetically distinct sub-Saharan African populations in the relatively distant past (Figure 8). The most recent pulse introduced FR ancestry 372, 95% CI [342, 418] generations ago, suggesting EA recently admixed with cosmopolitan populations, which is strikingly consistent with our estimates from the other admixed populations in Sub-Saharan Africa (Figure 3, Figure 8). Additionally, the relatively recent admixture between West African and Ethiopian *D. melanogaster*, suggests that Sub-Saharan populations of this species were isolated until very recently (i.e., approximately 338 years ago).

We note that our estimate of admixture time is congruent with an estimate of divergence between Ethiopian and Central African populations. Kern and Hey (2017) estimated Ethiopian and West African *D. melanogaster* diverged approximately 3628 years ago (or approximately 50,000 generations). Our estimates of admixture time occur sufficiently far after this estimated Ethiopian and African divergence time and provide confidence that our three-population admixture model of Gambella (EA) *D. melanogaster* is consistent with previous investigations of demographic patterns of *D. melanogaster* from sub-Saharan African. These data therefore indicate that our approach can be useful in estimating the timing of multiple admixture events in the history of a population using real genotype data.

**Figure 8.**
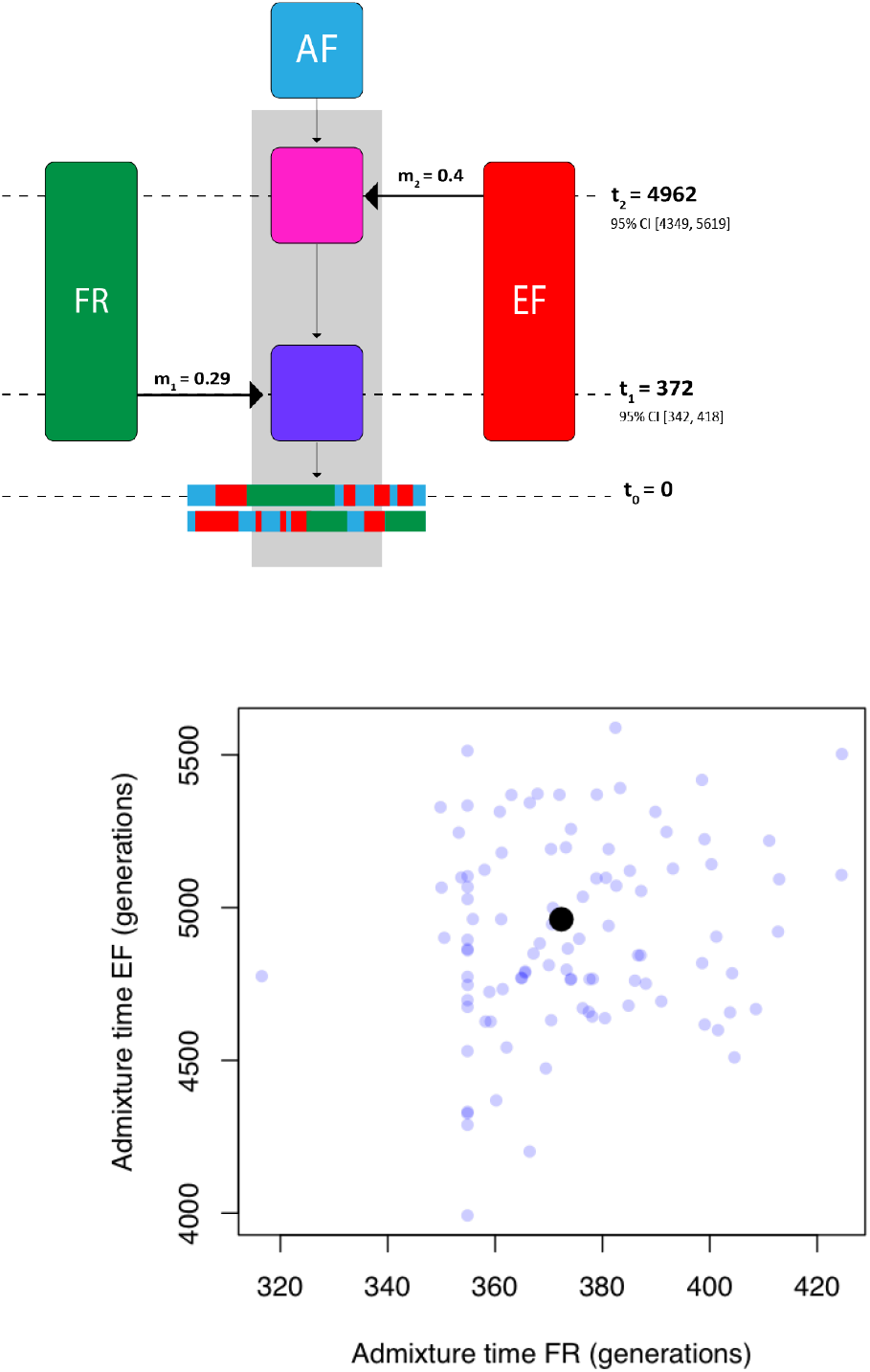
Most likely admixture model fitted to real data generated from Gambella, Ethiopia (EA). Schematic of double pulse admixture model with three ancestral populations and ancestry types is shown (top). Note the schematic is not drawn to scale. 1000 bootstraps were run of FR and EA pulsing into AF native ancestry. Real bootstrap data is shown (bottom), where each point is a bootstrap estimate of FR and EA admixture pulses and the black point represents the optimal admixture model from the full data set. The optima and 95% confidence intervals around the estimated timing of admixture are reported as part of the schematic.

## Conclusion

Admixture histories can be complex with numerous distinct ancestral populations contributing genetic material to a recipient population at multiple times in the past. In this work, we developed coalescent theory as well as a method to estimate the timing of multiple admixture events in an admixed population. We applied our model to forward-time simulated and real sequence data. The results of our simulations suggest that our model can discern between ancestry pulses occurring far apart in time from each other. Moreover, our method excels when the populations contributing ancestry are genetically diverse from each other.

Our approach, when applied to real admixed *Drosophila melanogaster* populations, is consistent with previous results on the African origin and admixture of the *Drosophila melanogaster* species. We find that cosmopolitan ancestry has entered very recently and is best accommodated by our framework using a single pulse model. Additionally, we demonstrate that more complex admixture patterns have shaped the ancestry of Gambella, Ethiopia. We indicate that two ancestry pulses from distinct ancestry types are necessary to explain patterns of genetic variation within this population.

This method improves the estimates of multiple pulse admixture models while accommodating the possibility of more than one source population. Although not a focus of this study, our method is designed to accommodate read pile-up data, samples of arbitrary ploidy, and samples with unknown admixture history. Taken all together, we anticipate that our approach will facilitate continued exploration of admixture’s contributions to fundamental biological processes such as adaptation, ecological divergence and speciation.

## Methods

### Coalescent Simulations

We simulated ancestral population genetic variation using the coalescent simulation software package MACS (Chen *et al*. 2009). Briefly, to consider a simplified ancestral process, we consider a model where a population, initially of size 2N, subdivided into three daughter populations of identical size, 2N. To assay a range of genetic divergence among the daughter populations, we performed replicate simulations where we allowed populations to diverge (D in the command line below) for 0.05, 0.1, 0.25, 0.5 and 1 N_*e*_ generations. Note that the functionally relevant parameter for LAI using this approach is probably related to allele frequency changes between the ancestral populations, and therefore to the density of ancestry informative markers along the genome, rather than to divergence times per s_*e*_. For example, a population bottleneck would result in rapid and dramatic allele frequency differences despite a short divergence time. Note also that alternative divergence models will also impact the distribution of LD among markers, underscoring the importance of considering each unique model when applying this approach for admixed model estimation.

We therefore used the following command line for all simulated populations:

`$ macs 600 100000000 -i 1 -h 100 -t 0.001 -r 0.001 -I 3 200 200 200 0 -ej D 2 1 -ej D 3 1`

This will simulate a single chromosome of length 100Mb with a per-site theta of 0.001 and equal recombination rate. Therefore, these simulations could resemble a reasonably sized mammalian population. This will output 200 chromosomes per ancestral population, and for all admixture simulations, we used the first 100 as the reference panel, and the second 100 to simulation genetic variation across admixed chromosomes.

### Admixture Simulations

We simulated admixed populations using the forward-in-time admixture simulation program, SELAM (Corbett-Detig and Jones 2016). We first explored the levels of LD pruning that is necessary for producing unbiased estimates of admixture times using two population, single pulse simulations. All simulations for LD pruning optimizations were run with ancestry proportions 0.5 and 0.5 and with an admixed population size of 5,000 males and 5,000 females.

We then simulated admixture models with three ancestral population models where the resident population contributed 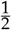 of the total ancestry, and each admixture pulse contributed 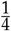th of the total ancestry to the final admixed population. Therefore the first pulse, in forward time, contributed 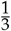 of the admixed population and then had 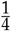th of that ancestry replaced by the subsequent pulse. All simulations were performed across the range of divergence times that we simulated for each ancestral population.

### Read Simulation

We simulated short read data for each admixed individual following the approach of (Corbett-Detig and Nielsen 2017). Briefly, reads are drawn binomially from each samples’ genotype and read depths from a Poisson distribution with mean equal to 2. That is, we simulated 2X sequencing depths to represent a likely use case of this software where individuals are sequenced to relatively light coverage.

### Drosophila Simulation

We have previously used this framework to simulate data consistent with the ancestral populations of *D. melanogaster*, following the coalescent simulation approach of (Corbett-Detig and Nielsen 2017). All other features of the simulated Drosophila dataset are similar to those above except that we simulated genotypes rather than short read pileup data. We did this because the dataset that we used is sequenced to sufficiently high depths so as to preclude most uncertainty in short read data (Lack *et al*. 2015, 2016).

### Robustness to Smaller Sample Sizes, Incorrect Recombination Rates, and Ancestral Population Divergence

We simulated varying sample sizes, erroneous recombination maps and reference population divergences to characterize our method’s ability to correctly estimate the timing of admixture. We simulated single admixture pulses 20, 40, 60, and 80 generations ago using sample sizes of 10, 25, and 50. We performed 20 replicate simulations. For all robustness checks, we used simulated data where the ancestral populations were 0.25 N_*e*_ generations divergent from one another.

To test for the effect of erroneous recombination maps, we modified the recombination map every 5Mb using the following approach. For each window, we do a draw of *D* from uniform(1 − d, 1 + d) where *d* is 0.25, 0.5, 0.75, or 1. We multiply D by the actual recombination rate at that window to obtain our modified recombination rate. For example, where *d* is 1, *D* is chosen from a uniform distribution of range(0,2). Where *d* is 0.25, *D* is chosen from a uniform distribution of range(0.75,1.25). Thus, the larger value of *d* the larger the recombination map will be perturbed.

We also tested our approach under scenarios where one of the reference populations used to estimate the timing of admixture is not the actual source population contributing to the admixed population. We estimated the timing of single and double pulse admixture events using a reference population that was 0.005, 0.01, 0.02, 0.05 and 0.1 N_*e*_ generations diverged from the real source population. We simulated single pulse data where the timing of admixture occurred 20, 40, 60, and 80 generations ago. We fit both single and double pulse admixture models to these data and computed summary statistics describing the fit of single pulse model to single pulse data. In addition, we simulated double pulse data where the timing of the most recent admixture pulse occurred 20, 40, 60, and 80 generations ago and where the most distant admixture pulse always occurred 100 generations ago. We computed summary statistics describing the fit of two pulse models to these data.

### Comparison to ALDER

We estimated the timing of single and double pulse admixtures using ALDER v1.02 (Loh *et al*. 2013) and MALDER v1.0 (*Pickrell *et al*.* 2014) respectively. We evaluated the results of our approach, ALDER, and MALDER by performing 20 replicates of each admixture scenario and computing the NRMSE for estimated admixture time and admixture proportion. We simulated single pulse admixture data using two reference populations of 50 individuals each and an admixed population of 50 individuals. Similarly, we simulated double pulse admixture data by adding a third reference population of 50 individuals. The timings of the single and double pulses are identical to those outlined above and can be found explicitly in Tables S13 and S14. ALDER and MALDER were run with a maxdis of 0.1. We note that the default settings of ALDER and MALDER could not detect admixture events occurring more than 100 generations ago. Thus, we extended the minimum distance (mindis) to 0 for any scenario with an admixture event occurring more than 100 generations ago. Extending the minimum distance to 0 may confound estimated admixture times because ancestral linkage disequilibrium will have a greater impact on short scales, but appeared to produce better admixture time estimates for the scenarios we considered. We note that there may be other ways to improve ALDER and MALDER’s performance in detecting ancient admixture events.

### Drosophila Sample Application

We obtained *Drosophila melanogaster* genotype data for natural isolates from the Drosophila Genome Nexus (Lack *et al*. 2015, 2016), which curates data from more than 1,000 natural isolates of this species. We used the recommended masking packages supplied for that site for all samples in reference populations. For SD, we used the ZI population, for RG we used a set of central and west African samples (as in Corbett-Detig and Nielsen (2017)), and for EA we used the EF population. All cosmopolitan pulses were modeled using FR as the cosmopolitan reference population.

### Application to Admixed Samples

We used our model to assay the admixture histories of Rwandan, South African, and Ethiopian samples. We fit both a single pulse model and double pulse model to genotype data from each population, running 100 bootstrap replicates using a block size of 5000 SNPs.

`- single pulse`

`./ancestry_hmm -i RG.auto.panel -s RG.ploidy.txt -a 2 0.1 0.9 -p 1 100000 0.9 -p 0 -100 0.1 -g -e 1e-4 -b 100 ↪ 5000`

`./ancestry_hmm -i SD.auto.panel -s SD.ploidy.txt -a 2 0.17 0.83 -p 1 100000 0.83 -p 0 -100 0.17 -g -e 1e-4 -b ↪ 100 5000`

`./ancestry_hmm -i EA.auto.panel -s EA.ploidy.fixed.txt -a 2 0.34 0.66 -p 1 100000 0.66 -p 0 -100 0.34 -g -e ↪ 1e-4 -b 100 5000`

`- double pulse`

`./ancestry_hmm -i RG.auto.panel -s RG.ploidy.txt -a 2 0.1 0.9 -p 1 100000 0.9 -p 0 -100 -0.05 -p 0 -100 -0.05 ↪ -e 1e-4 -g -b 100 5000`

`./ancestry_hmm -i SD.auto.panel -s SD.ploidy.txt -a 2 0.17 0.83 -p 1 100000 0.83 -p 0 -100 -0.085 -p 0 -100 ↪ -0.085 -e 1e-4 -g -b 100 5000`

`./ancestry_hmm -i EA.auto.panel -s EA.ploidy.txt -a 2 0.34 0.66 -p 1 100000 0.66 -p 0 -100 -0.17 -p 0 -100 ↪ -0.17 -e 1e-4 -g -b 100 5000`

### Application to Three Population Mixture from Gambella, Ethiopia

We used our model to assay the admixture histories of Gambella, Ethiopia. We fit a double pulse model to genotype data from Gambella, running 1000 bootstrap replicates each using a block size of 5000 SNPs. Moreover, since the native ancestry type for Gambella is unknown, we modeled West African (AF), Ethiopian (EA), and French (FR) as the ancestral population.

`- EA native ancestry`

`./ancestry_hmm -i three_pop.auto.EA.panel -s three_pop.auto.EA.ploidy -a 3 0.4 0.31 0.29 -p 0 -1000 0.4 -p 1 ↪ 100000 0.31 -p 2 -1000 0.29 -g -b 1000 5000`

## Supplemental Material

**Table S1.** Description of summary statistics used to describe admixture models and LAI accuracy.

| Label | Description |
| --- | --- |
| pulse_time | True admixture time simulated using SELAM. |
| Normalized root mean squared error | across 20 replicate simulations. We normalized each error estimate by the true admixture time or ancestry proportion. |
| mean_posterior_error | The average distance between the posterior distribution of hidden states and the true state. See Corbett-Detig and Nielsen (2017) for a formal explanation. |
| accuracy | The proportion of sites where the maximum likelihood estimate of the hidden state is equal to the true ancestry state after LAI. |
| confident_correct | The proportion of accurate sites where the posterior probability is greater than 0.95 after LAI. |

**Supplementary Figure S1.**
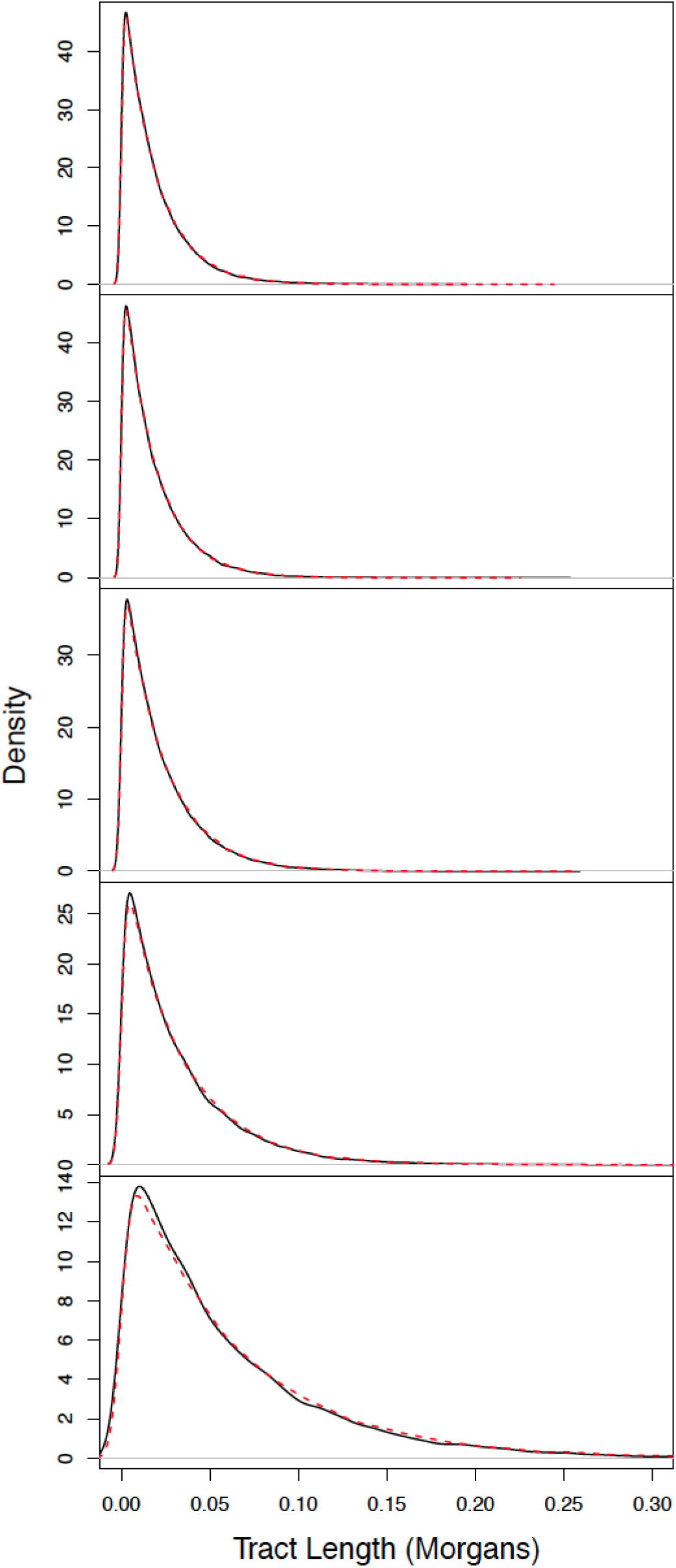
Tract length distributions obtained using our tract length model approximation (solid black) and forward-in-time simulation (dashed red). In the model considered, there are five ancestry types *A*_1_‥*A_k_* and four admixture pulses occurring at 20, 40, 60, and 80 generations since the present. Each pulse in forward time contributes 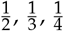, and 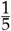, respectively, of total resident genetic ancestry. Each simulation had a diploid population size of size 10,000 and we aggregated data from 50 sampled individuals across 100 simulations to produce the full tract length distribution. From top to bottom, respective ancestry tract length distributions correspond *A_k_*, *A*_1_, *A*_2_, *A*_3_, *A*_4_.

**Table S2.** Simulated and analytical transition rates for a model with five ancestry types, *k* = 5. *t*_4_, *t*_3_, *t*_2_, *t*_1_, and *t*_0_ are 80, 60,40, 20, and 0 generations since the present, respectively. *m*_4_, *m*_3_, *m*_2_, and *m*_1_ are 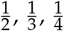, and 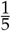 respectively.

| From | To | Rate_Simulation | Rate_Analytical |
| --- | --- | --- | --- |
| 5 | 4 | 25.715 | 25.625 |
| 5 | 3 | 15.509 | 15.645 |
| 5 | 2 | 9.045 | 8.991 |
| 5 | 1 | 4.025 | 3.998 |
| 4 | 5 | 25.449 | 25.625 |
| 4 | 3 | 15.654 | 15.645 |
| 4 | 2 | 8.956 | 8.991 |
| 4 | 1 | 3.978 | 3.998 |
| 3 | 5 | 15.640 | 15.645 |
| 3 | 4 | 15.509 | 15.645 |
| 3 | 2 | 8.920 | 8.991 |
| 3 | 1 | 4.013 | 3.998 |
| 2 | 5 | 9.007 | 8.991 |
| 2 | 4 | 9.066 | 8.991 |
| 2 | 3 | 9.058 | 8.991 |
| 2 | 1 | 3.955 | 3.998 |
| 1 | 5 | 3.936 | 3.998 |
| 1 | 4 | 4.001 | 3.998 |
| 1 | 3 | 3.970 | 3.998 |
| 1 | 2 | 4.001 | 3.998 |

**Supplementary Figure S2.**
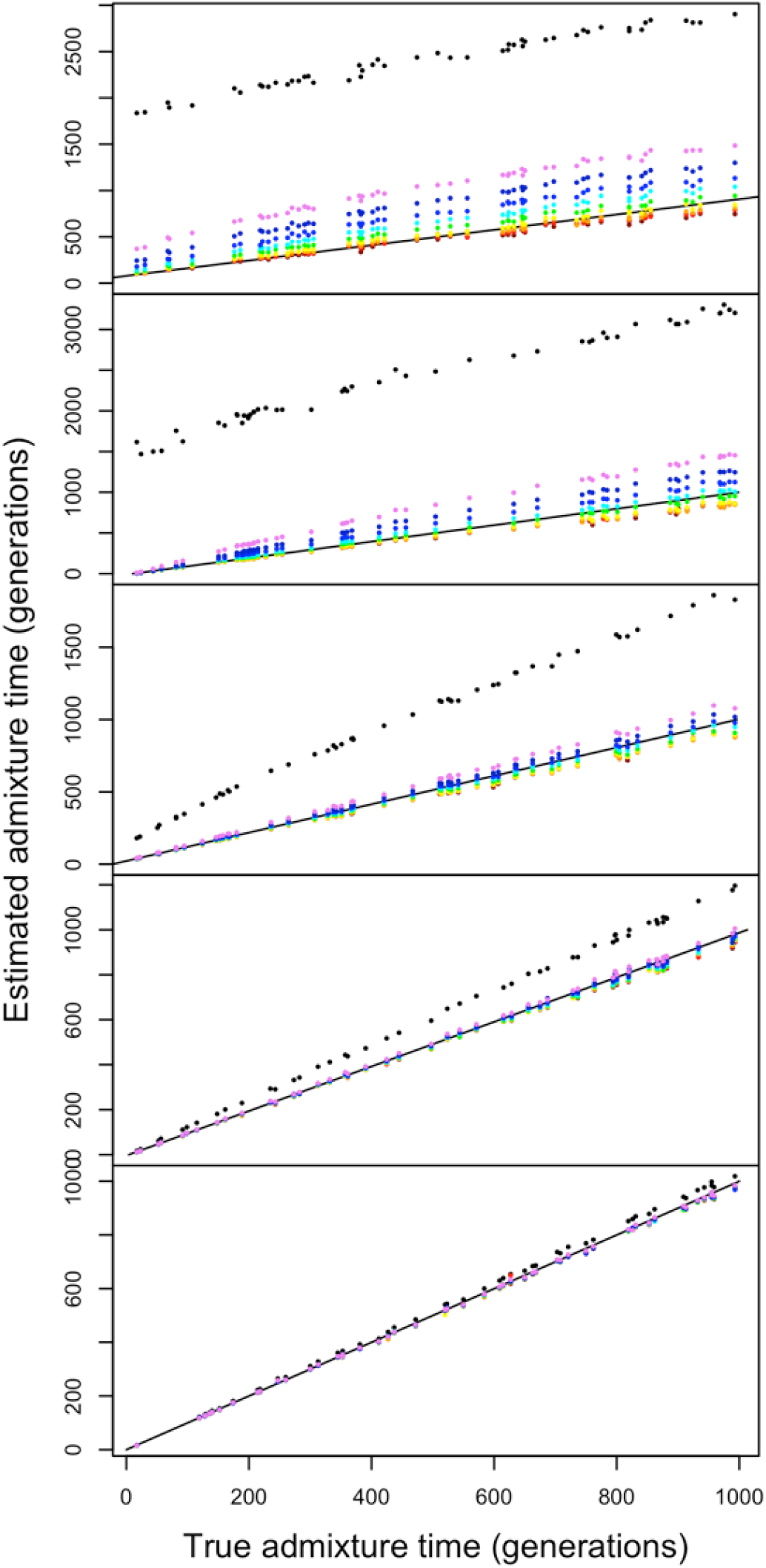
Estimated admixture times for single pulse admixture models with varying levels of genetic divergence between ancestral populations. Specifically, from top to bottom, ancestral populations are 0.05, 0.1, 0.25, 0.5, and 1 N_*e*_ generations divergent from one another. LD pruned is 1 (black), 0.9 (violet), 0.8 (dark blue), 0.7 (blue), 0.6 (cyan), 0.5 (green), 0.4 (yellow), 0.3 (orange), 0.2 (red), 0.1 (dark red).

**Supplementary Figure S3.**
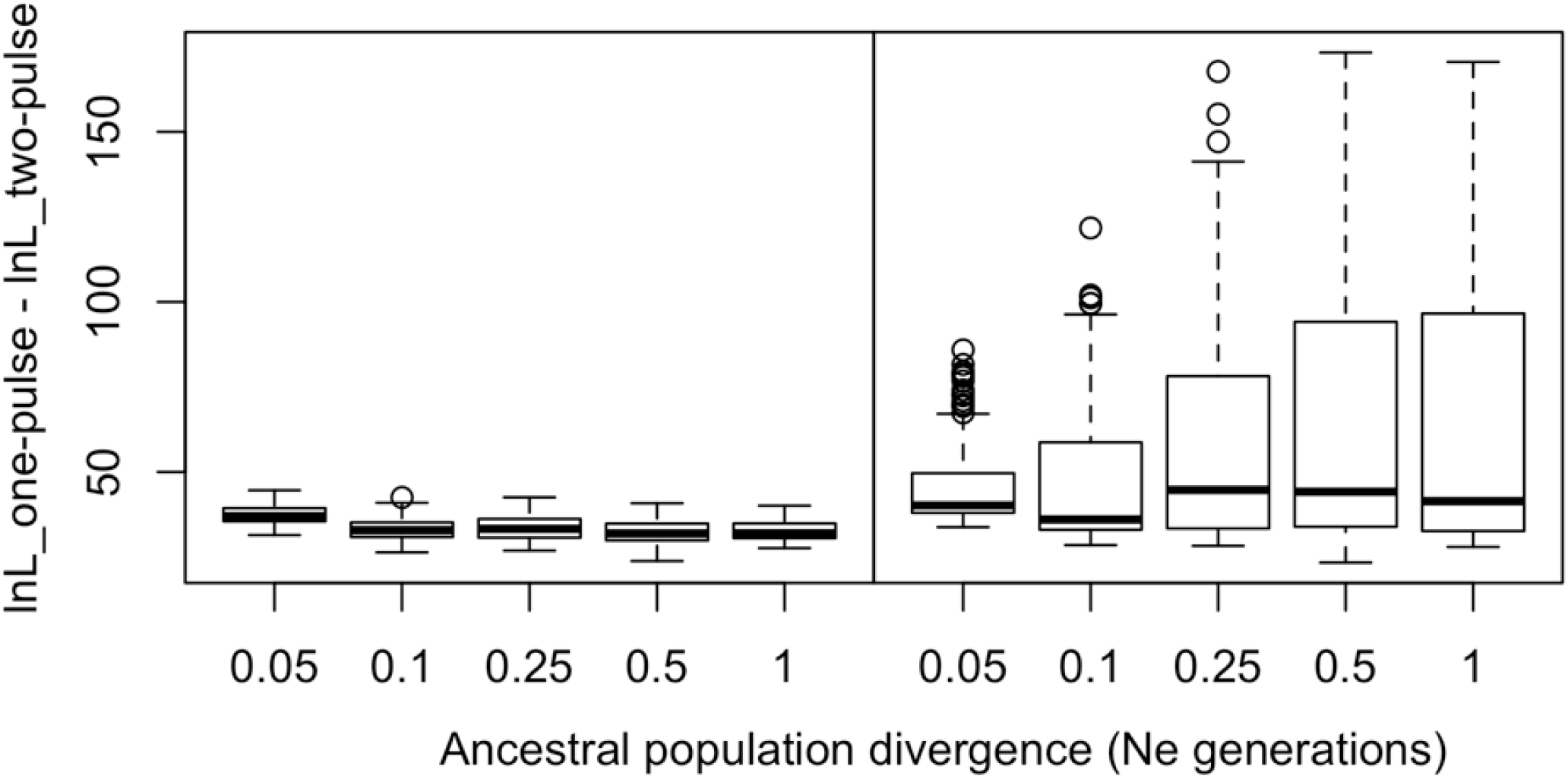
The likelihood ratio between two-pulse and single-pulse ancestry models fit to data simulated under a single pulse (left) and a two pulse model (right). The single pulse model included a pulse of 50% of ancestry 50 generations prior to sampling. The two pulse model included a first pulse in forward time of 33% of ancestry 100 generations prior to sampling, and a second pulse in forward time of 25% of ancestry both from ancestry type 1, at time 20 generations prior to sampling. Each boxplot is based on 20 replicate simulations for each level of ancestral population divergence.

**Supplementary Figure S4.**
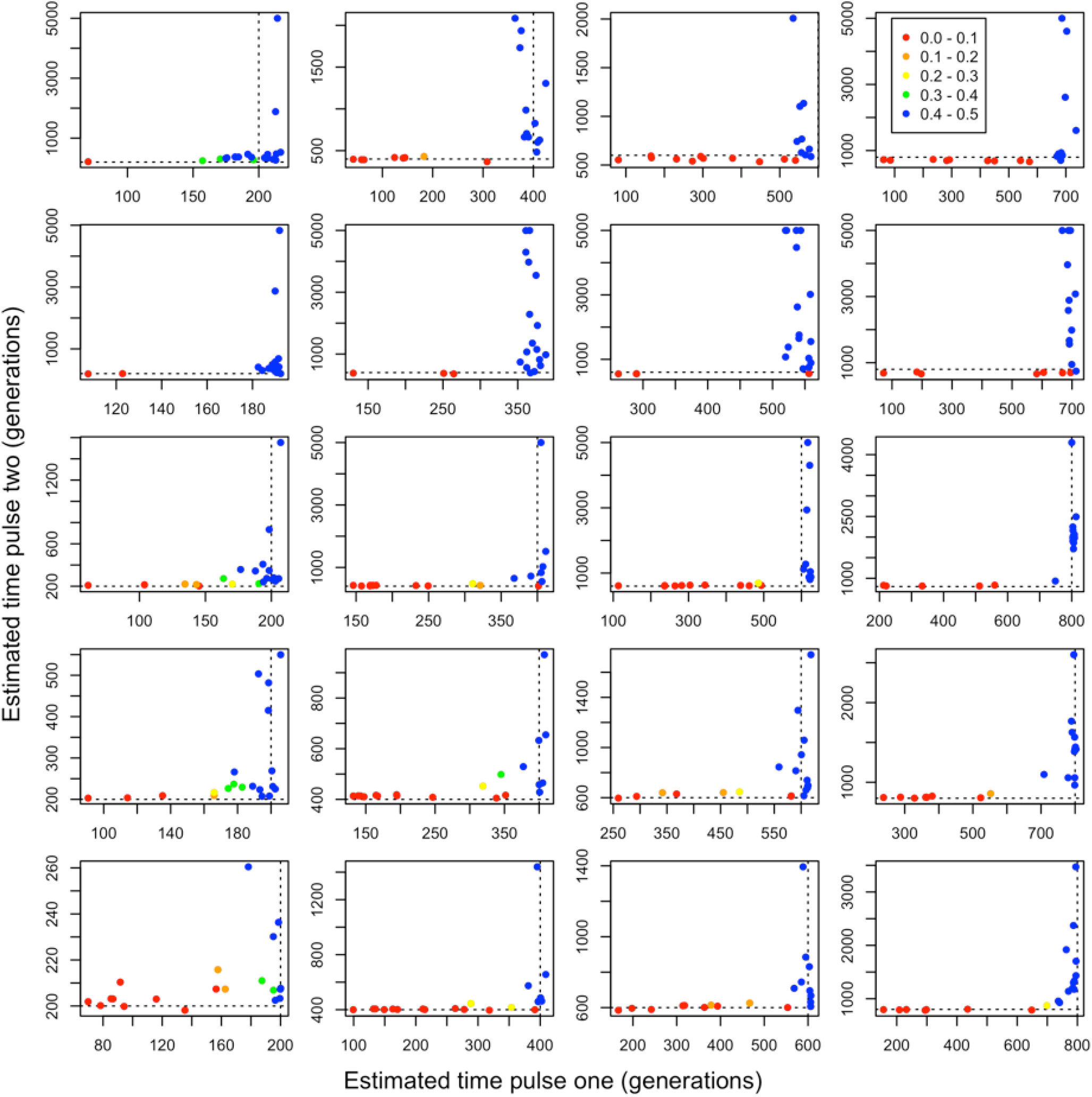
Two-pulse admixture models fitted using our framework to data generated under a single pulse admixture model. We considered varying levels of population divergence. From top to bottom, the ancestral populations are 0.05, 0.1, 0.25, 0.5, and 1 N_*e*_ generation divergent from one another. From left to right, the single admixture pulse occurred 200, 400, 600, and 800 generations prior to sampling and replaced one half of the individuals in the population. Point colors correspond to the proportion of ancestry that is attributable to the second pulse with 0.0-0.1 (red), 0.1-0.2 (orange), 0.2-0.3 (yellow), 0.3-0.4 (green) and 0.4-0.5 (blue). Dashed lines reflect the true admixture time. Note that axes may differ between subplots.

**Table S3.** Local ancestry inference accuracy statistics of single pulse admixture models fit to two pulse admixture data. Listed statistics are described in Table S1.

| divergence | pulse_time | mean_posterior_error | accuracy | confident_correct |
| --- | --- | --- | --- | --- |
| 0.05 | 20 | 0.0086 | 0.7991 | 0.2878 |
| 0.05 | 40 | 0.0071 | 0.7837 | 0.2383 |
| 0.05 | 60 | 0.0060 | 0.7698 | 0.1981 |
| 0.05 | 80 | 0.0053 | 0.7582 | 0.1712 |
| 0.05 | 200 | 0.0004 | 0.5092 | 0.0037 |
| 0.05 | 400 | 0.0002 | 0.4903 | 0.0023 |
| 0.05 | 600 | 0.0002 | 0.4751 | 0.001 |
| 0.05 | 800 | 0.0001 | 0.4659 | 0.0016 |
| 0.1 | 20 | 0.0080 | 0.9331 | 0.7713 |
| 0.1 | 40 | 0.0079 | 0.9278 | 0.7471 |
| 0.1 | 60 | 0.0078 | 0.9216 | 0.7221 |
| 0.1 | 80 | 0.0080 | 0.91638 | 0.7019 |
| 0.1 | 200 | 0.0046 | 0.6983 | 0.0999 |
| 0.1 | 400 | 0.0030 | 0.676 | 0.0675 |
| 0.1 | 600 | 0.0025 | 0.6621 | 0.0525 |
| 0.1 | 800 | 0.0021 | 0.6504 | 0.0437 |
| 0.25 | 20 | 0.0010 | 1 | 0.9697 |
| 0.25 | 40 | 0.0011 | 0.9902 | 0.9669 |
| 0.25 | 60 | 0.001 | 0.9892 | 0.9634 |
| 0.25 | 80 | 0.0013 | 0.9884 | 0.9606 |
| 0.25 | 200 | 0.0093 | 0.9174 | 0.7203 |
| 0.25 | 400 | 0.0093 | 0.9106 | 0.6935 |
| 0.25 | 600 | 0.0092 | 0.9039 | 0.6667 |
| 0.25 | 800 | 0.0094 | 0.8979 | 0.6447 |
| 0.5 | 20 | 0.0002 | 0.9974 | 0.9912 |
| 0.5 | 40 | 0.0002 | 0.9972 | 0.9904 |
| 0.5 | 60 | 0.0003 | 0.9969 | 0.9894 |
| 0.5 | 80 | 0.0003 | 0.9967 | 0.98873 |
| 0.5 | 200 | 0.0035 | 0.9724 | 0.9088 |
| 0.5 | 400 | 0.0037 | 0.9696 | 0.8989 |
| 0.5 | 600 | 0.0039 | 0.9673 | 0.8908 |
| 0.5 | 800 | 0.0041 | 0.9649 | 0.8825 |
| 1 | 20 | 6.0757e-05 | 0.9993 | 0.99764 |
| 1 | 40 | 6.7744e-05 | 0.9993 | 0.9974 |
| 1 | 60 | 7.4457e-05 | 0.99916 | 0.99720 |
| 1 | 80 | 8.0574e-05 | 0.9991 | 0.9969 |
| 1 | 200 | 0.0008 | 0.9925 | 0.97535 |
| 1 | 400 | 0.0009 | 0.9918 | 0.97289 |
| 1 | 600 | 0.0010 | 0.9909 | 0.97002 |
| 1 | 800 | 0.0011 | 0.9903 | 0.96794 |

**Table S4.** Local ancestry inference accuracy statistics of two pulse admixture model fit to two pulse admixture population. Listed statistics are described in Table S1.

| divergence | pulse_time | mean_posterior_error | accuracy | confident_correct |
| --- | --- | --- | --- | --- |
| 0.05 | 20 | 0.0067 | 0.7978 | 0.2591 |
| 0.05 | 40 | 0.0062 | 0.7825 | 0.2211 |
| 0.05 | 60 | 0.0054 | 0.7686 | 0.1873 |
| 0.05 | 80 | 0.0049 | 0.7572 | 0.1631 |
| 0.05 | 200 | 0.0003 | 0.5093 | 0.0034 |
| 0.05 | 400 | 0.0002 | 0.4902 | 0.0023 |
| 0.05 | 600 | 0.0002 | 0.4753 | 0.0018 |
| 0.05 | 800 | 0.0001 | 0.466 | 0.0016 |
| 0.1 | 20 | 0.0074 | 0.9333 | 0.7648 |
| 0.1 | 40 | 0.0077 | 0.9277 | 0.7434 |
| 0.1 | 60 | 0.0078 | 0.9216 | 0.7215 |
| 0.1 | 80 | 0.0080 | 0.9164 | 0.7016 |
| 0.1 | 200 | 0.0037 | 0.6979 | 0.0898 |
| 0.1 | 400 | 0.0029 | 0.6759 | 0.0661 |
| 0.1 | 600 | 0.0025 | 0.662 | 0.0525 |
| 0.1 | 800 | 0.0021 | 0.6505 | 0.0438 |
| 0.25 | 20 | 0.0010 | 0.991 | 0.9693 |
| 0.25 | 40 | 0.0011 | 0.9902 | 0.9666 |
| 0.25 | 60 | 0.0012 | 0.9892 | 0.9633 |
| 0.25 | 80 | 0.0013 | 0.9884 | 0.9604 |
| 0.25 | 200 | 0.0085 | 0.9175 | 0.7112 |
| 0.25 | 400 | 0.0090 | 0.9105 | 0.6882 |
| 0.25 | 600 | 0.0091 | 0.9039 | 0.6648 |
| 0.25 | 800 | 0.0094 | 0.8979 | 0.6440 |
| 0.5 | 20 | 0.0002 | 0.9974 | 0.9911 |
| 0.5 | 40 | 0.0002 | 0.9972 | 0.9904 |
| 0.5 | 60 | 0.0003 | 0.9969 | 0.9893 |
| 0.5 | 80 | 0.0003 | 0.9967 | 0.9886 |
| 0.5 | 200 | 0.0033 | 0.9724 | 0.9071 |
| 0.5 | 400 | 0.0036 | 0.9696 | 0.8978 |
| 0.5 | 600 | 0.0039 | 0.9673 | 0.8903 |
| 0.5 | 800 | 0.0041 | 0.9648 | 0.8823 |
| 1 | 20 | 6.0073e-05 | 0.9993 | 0.9976 |
| 1 | 40 | 6.7661e-05 | 0.9993 | 0.9974 |
| 1 | 60 | 7.4321e-05 | 0.9992 | 0.9972 |
| 1 | 80 | 8.0529e-05 | 0.9991 | 0.9969 |
| 1 | 200 | 0.0008 | 0.9925 | 0.9752 |
| 1 | 400 | 0.0009 | 0.9918 | 0.9727 |
| 1 | 600 | 0.0010 | 0.9909 | 0.9698 |
| 1 | 800 | 0.0011 | 0.9903 | 0.9679 |

**Table S5.**
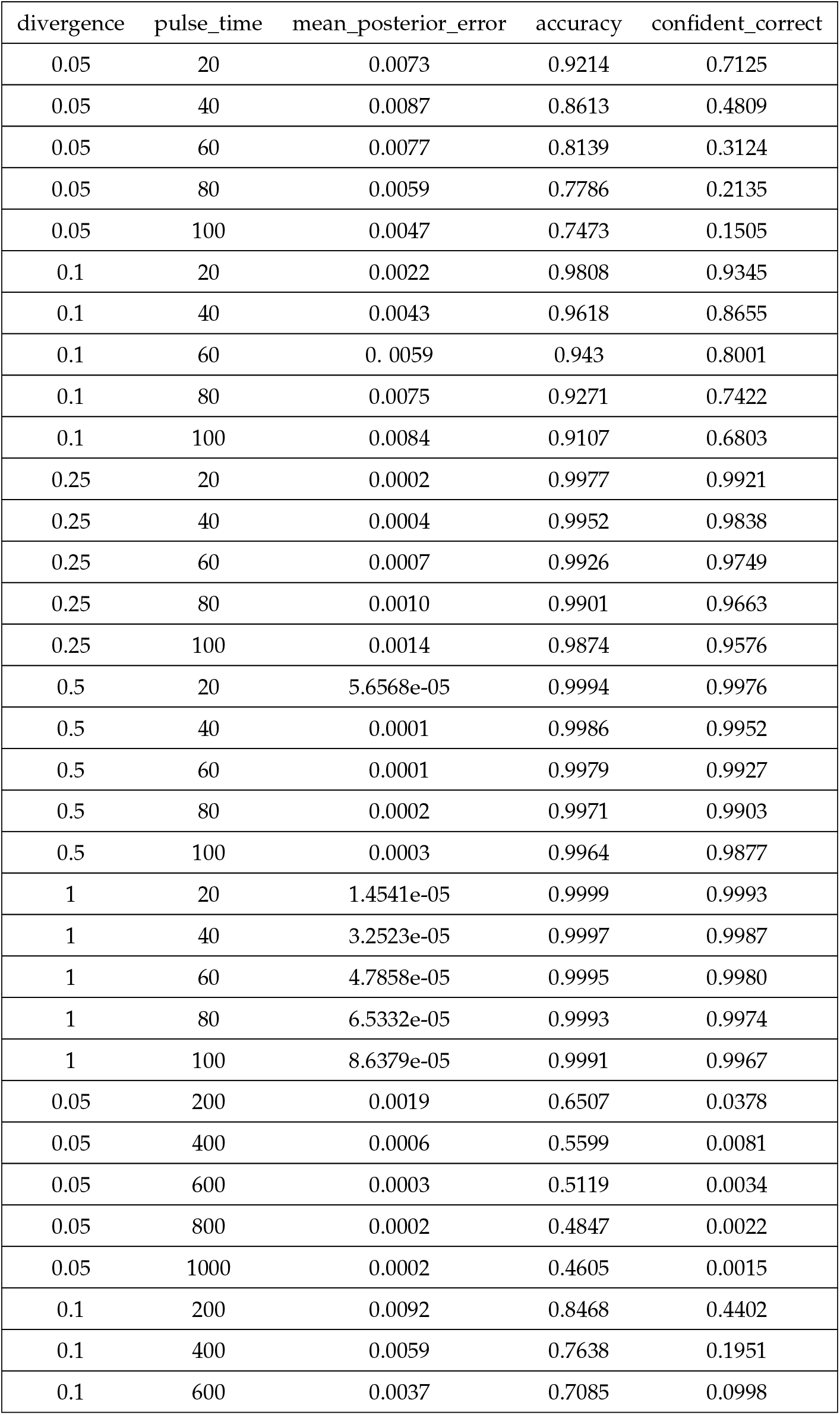
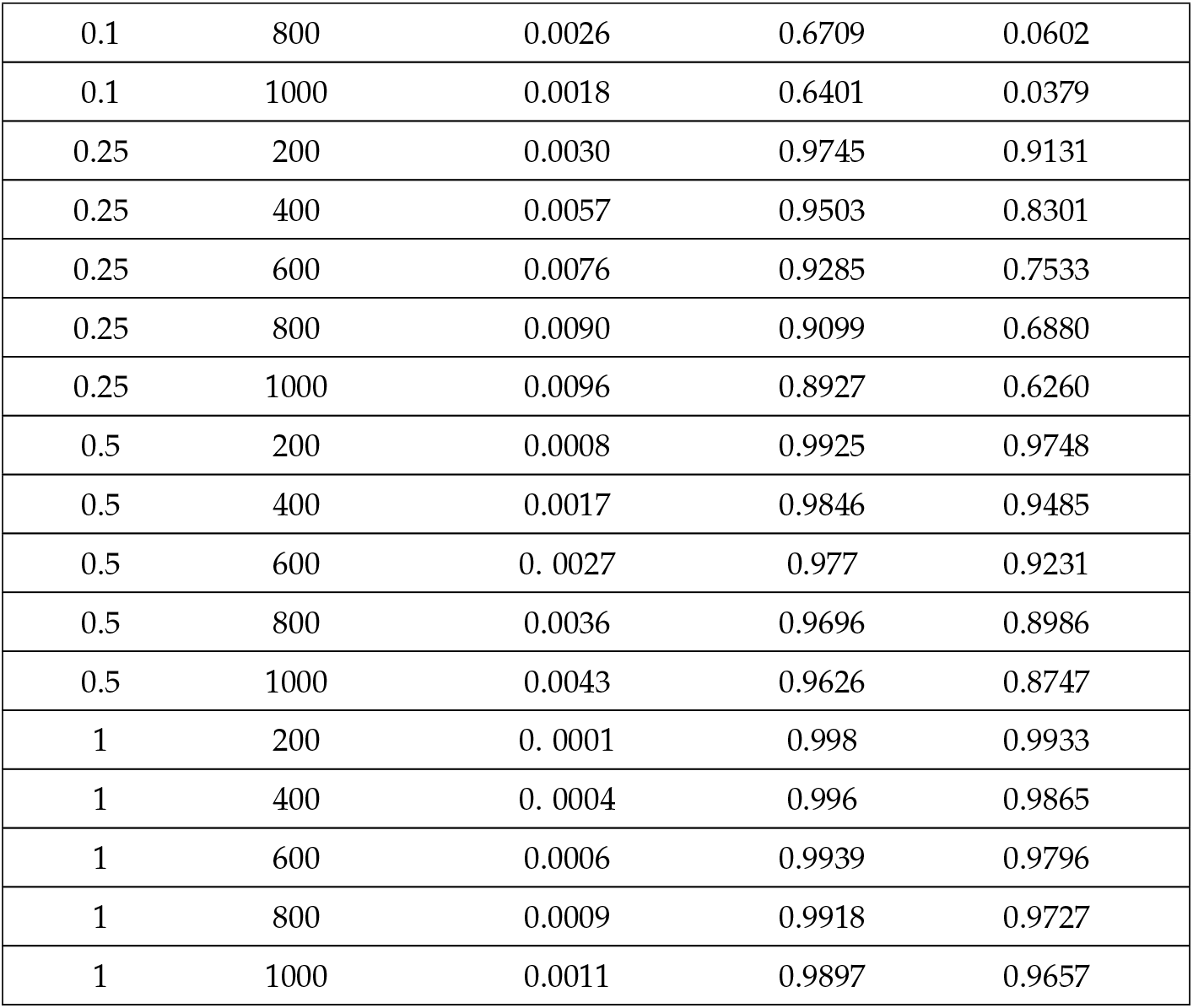
Local ancestry inference accuracy statistics of two pulse admixture model fit to single pulse admixture population. Listed statistics are described in Table S1.

| divergence | pulse_time | mean_posterior_error | accuracy | confident_correct |
| --- | --- | --- | --- | --- |
| 0.05 | 20 | 0.0073 | 0.9214 | 0.7125 |
| 0.05 | 40 | 0.0087 | 0.8613 | 0.4809 |
| 0.05 | 60 | 0.0077 | 0.8139 | 0.3124 |
| 0.05 | 80 | 0.0059 | 0.7786 | 0.2135 |
| 0.05 | 100 | 0.0047 | 0.7473 | 0.1505 |
| 0.1 | 20 | 0.0022 | 0.9808 | 0.9345 |
| 0.1 | 40 | 0.0043 | 0.9618 | 0.8655 |
| 0.1 | 60 | 0.0059 | 0.943 | 0.8001 |
| 0.1 | 80 | 0.0075 | 0.9271 | 0.7422 |
| 0.1 | 100 | 0.0084 | 0.9107 | 0.6803 |
| 0.25 | 20 | 0.0002 | 0.9977 | 0.9921 |
| 0.25 | 40 | 0.0004 | 0.9952 | 0.9838 |
| 0.25 | 60 | 0.0007 | 0.9926 | 0.9749 |
| 0.25 | 80 | 0.0010 | 0.9901 | 0.9663 |
| 0.25 | 100 | 0.0014 | 0.9874 | 0.9576 |
| 0.5 | 20 | 5.6568e-05 | 0.9994 | 0.9976 |
| 0.5 | 40 | 0.0001 | 0.9986 | 0.9952 |
| 0.5 | 60 | 0.0001 | 0.9979 | 0.9927 |
| 0.5 | 80 | 0.0002 | 0.9971 | 0.9903 |
| 0.5 | 100 | 0.0003 | 0.9964 | 0.9877 |
| 1 | 20 | 1.4541e-05 | 0.9999 | 0.9993 |
| 1 | 40 | 3.2523e-05 | 0.9997 | 0.9987 |
| 1 | 60 | 4.7858e-05 | 0.9995 | 0.9980 |
| 1 | 80 | 6.5332e-05 | 0.9993 | 0.9974 |
| 1 | 100 | 8.6379e-05 | 0.9991 | 0.9967 |
| 0.05 | 200 | 0.0019 | 0.6507 | 0.0378 |
| 0.05 | 400 | 0.0006 | 0.5599 | 0.0081 |
| 0.05 | 600 | 0.0003 | 0.5119 | 0.0034 |
| 0.05 | 800 | 0.0002 | 0.4847 | 0.0022 |
| 0.05 | 1000 | 0.0002 | 0.4605 | 0.0015 |
| 0.1 | 200 | 0.0092 | 0.8468 | 0.4402 |
| 0.1 | 400 | 0.0059 | 0.7638 | 0.1951 |
| 0.1 | 600 | 0.0037 | 0.7085 | 0.0998 |
| 0.1 | 800 | 0.0026 | 0.6709 | 0.0602 |
| 0.1 | 1000 | 0.0018 | 0.6401 | 0.0379 |
| 0.25 | 200 | 0.0030 | 0.9745 | 0.9131 |
| 0.25 | 400 | 0.0057 | 0.9503 | 0.8301 |
| 0.25 | 600 | 0.0076 | 0.9285 | 0.7533 |
| 0.25 | 800 | 0.0090 | 0.9099 | 0.6880 |
| 0.25 | 1000 | 0.0096 | 0.8927 | 0.6260 |
| 0.5 | 200 | 0.0008 | 0.9925 | 0.9748 |
| 0.5 | 400 | 0.0017 | 0.9846 | 0.9485 |
| 0.5 | 600 | 0.0027 | 0.977 | 0.9231 |
| 0.5 | 800 | 0.0036 | 0.9696 | 0.8986 |
| 0.5 | 1000 | 0.0043 | 0.9626 | 0.8747 |
| 1 | 200 | 0.0001 | 0.998 | 0.9933 |
| 1 | 400 | 0.0004 | 0.996 | 0.9865 |
| 1 | 600 | 0.0006 | 0.9939 | 0.9796 |
| 1 | 800 | 0.0009 | 0.9918 | 0.9727 |
| 1 | 1000 | 0.0011 | 0.9897 | 0.9657 |

**Supplementary Figure S5.**
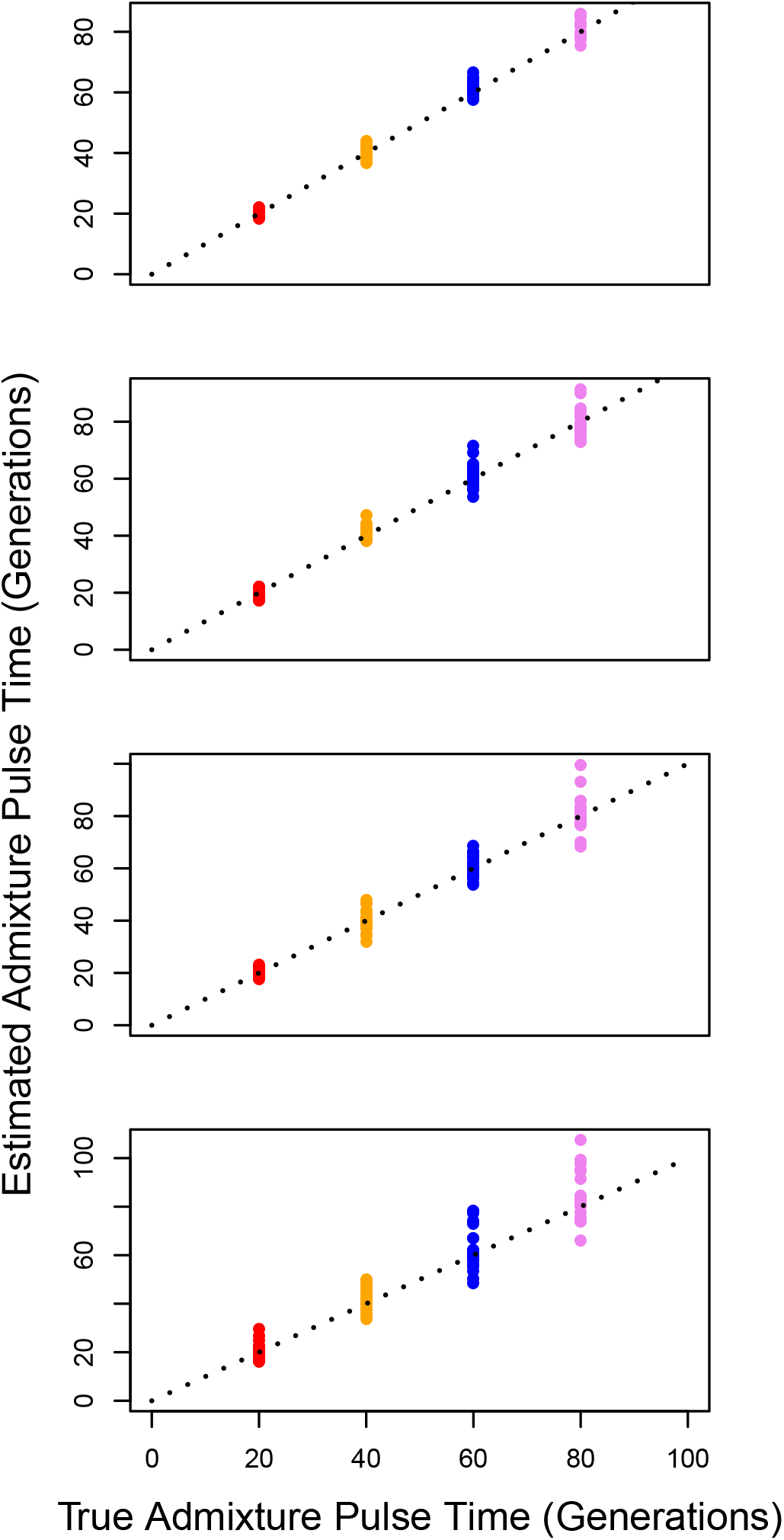
Admixture time estimates for single pulse models fitted to single pulse data at varying levels of recombination map perturbation. Rho was modified every 5Mb by multiplying by a scalar chosen at random from a uniform distribution between 1-d and 1+d where d is 0.25, 0.5 0.75 and 1 from top to bottom (See Methods). We simulated single pulse models occurring 20 (red), 40 (orange), 60 (blue), and 80 (pink) generations ago. Note that axes may differ between subplots. Summary statistics of model fitting are reported in Tables S6 and S7.

**Table S6.** Accuracy statistics of estimated single pulse admixture time fit to data generated under a single pulse model at varying levels of recombination map perturbation. Listed statistics are described in Table S1.

| pulse_time | NRMSE_0.25 | NRMSE_0.5 | NRMSE_0.75 | NRMSE_1.0 |
| --- | --- | --- | --- | --- |
| 20 | 0.0314 | 0.0522 | 0.0805 | 0.1169 |
| 40 | 0.0400 | 0.0534 | 0.0739 | 0.1153 |
| 60 | 0.0401 | 0.0594 | 0.0588 | 0.1186 |
| 80 | 0.0327 | 0.0554 | 0.0619 | 0.1014 |

**Table S7.** Local ancestry inference accuracy statistics of single pulse admixture model fit to single pulse admixture population after perturbing recombination rates every 5Mb windowed. Listed statistics are described in Table S1.

| pulse_time | rho_error | mean_posterior_error | accuracy | confident_correct |
| --- | --- | --- | --- | --- |
| 20 | 0.25 | 0.0089 | 0.9890 | 0.9837 |
| 20 | 0.5 | 0.0089 | 0.9890 | 0.9837 |
| 20 | 0.75 | 0.0089 | 0.9890 | 0.9837 |
| 20 | 1 | 0.0089 | 0.9889 | 0.9836 |
| 40 | 0.25 | 0.0140 | 0.9816 | 0.9707 |
| 40 | 0.5 | 0.0141 | 0.9816 | 0.9707 |
| 40 | 0.75 | 0.0141 | 0.9816 | 0.9706 |
| 40 | 1 | 0.0141 | 0.9816 | 0.9704 |
| 60 | 0.25 | 0.0180 | 0.9755 | 0.9588 |
| 60 | 0.5 | 0.0180 | 0.9755 | 0.9587 |
| 60 | 0.75 | 0.0181 | 0.9754 | 0.9584 |
| 60 | 1 | 0.0181 | 0.9754 | 0.9583 |
| 80 | 0.25 | 0.0204 | 0.9710 | 0.9485 |
| 80 | 0.5 | 0.0204 | 0.9710 | 0.9484 |
| 80 | 0.75 | 0.0204 | 0.9709 | 0.9482 |
| 80 | 1 | 0.0205 | 0.9708 | 0.9478 |

**Supplementary Figure S6.**
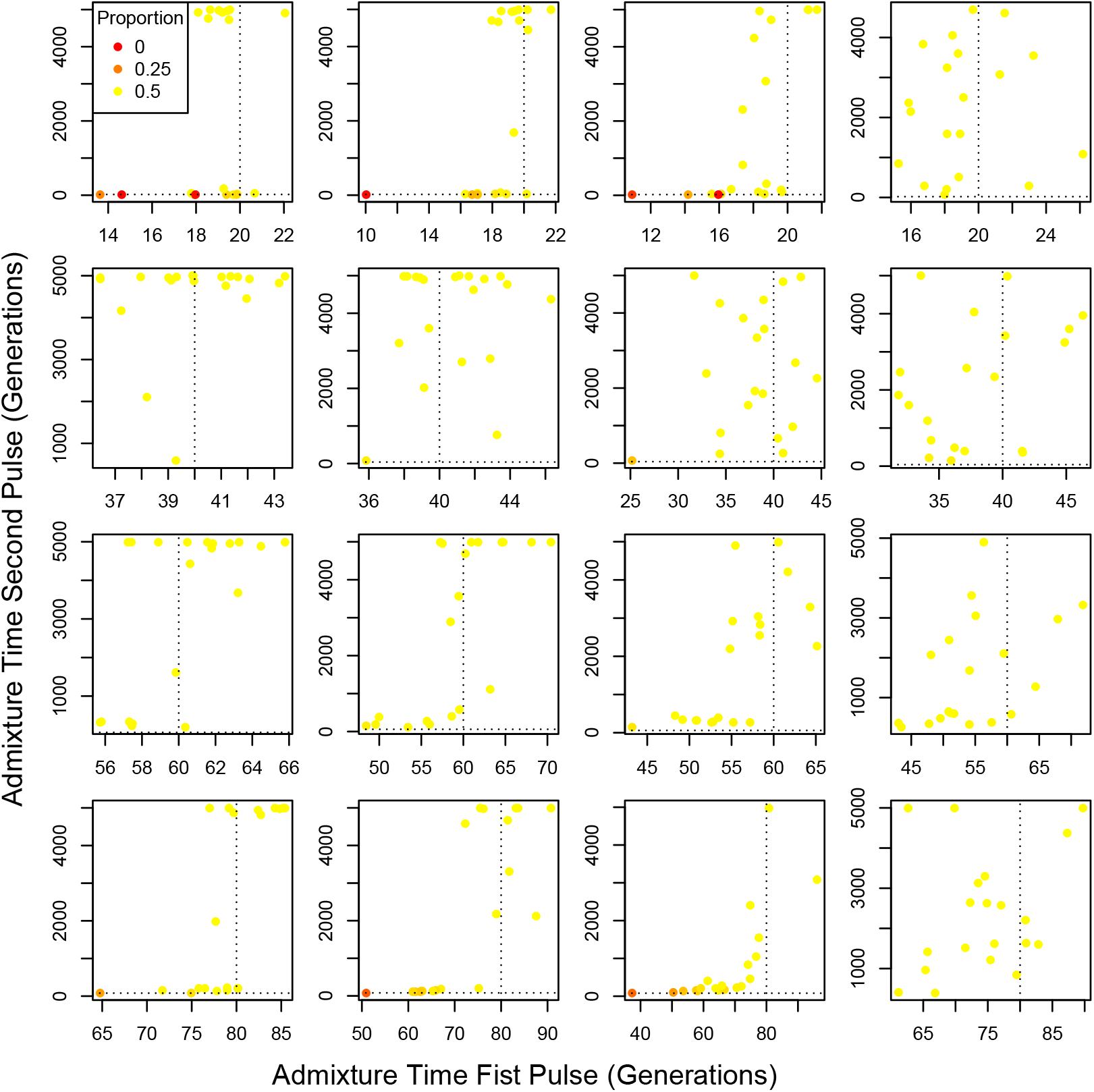
Admixture time estimates for double pulse models fitted to single pulse data at varying levels of recombination map perturbation. Rho was modified every 5Mb by multiplying by a scalar chosen at random from a uniform distribution between 1-d and 1+d where d is 0.25, 0.5 0.75 and 1 from left to right. For all admixture models considered, the second pulse occurred 100 generations before the present. From top to bottom, the most recent pulse occurred 20, 40, 60, and 80 generations ago. Note that axes may differ between subplots.

**Supplementary Figure S7.**
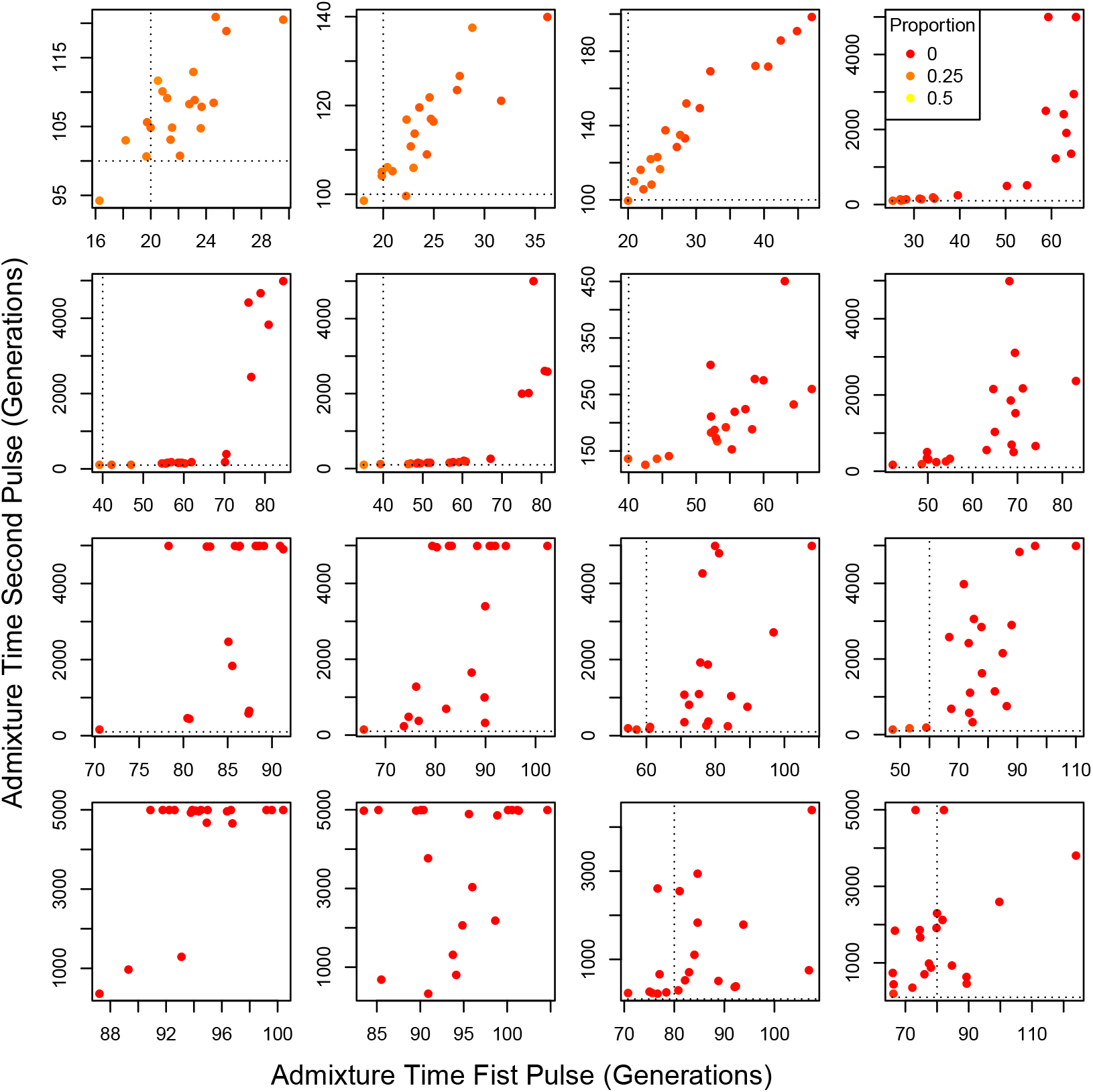
Admixture time estimates for double pulse models fitted to double pulse data at varying levels of recombination map perturbation. Rho was modified every 5Mb by multiplying by a scalar chosen at random from a uniform distribution between 1-d and 1+d where d is 0.25, 0.5 0.75 and 1 from left to right. For all admixture models considered, the second pulse occurred 100 generations before the present. From top to bottom, the most recent pulse occurred 20, 40, 60, and 80 generations ago. Note that axes may differ between subplots.

**Supplementary Figure S8.**
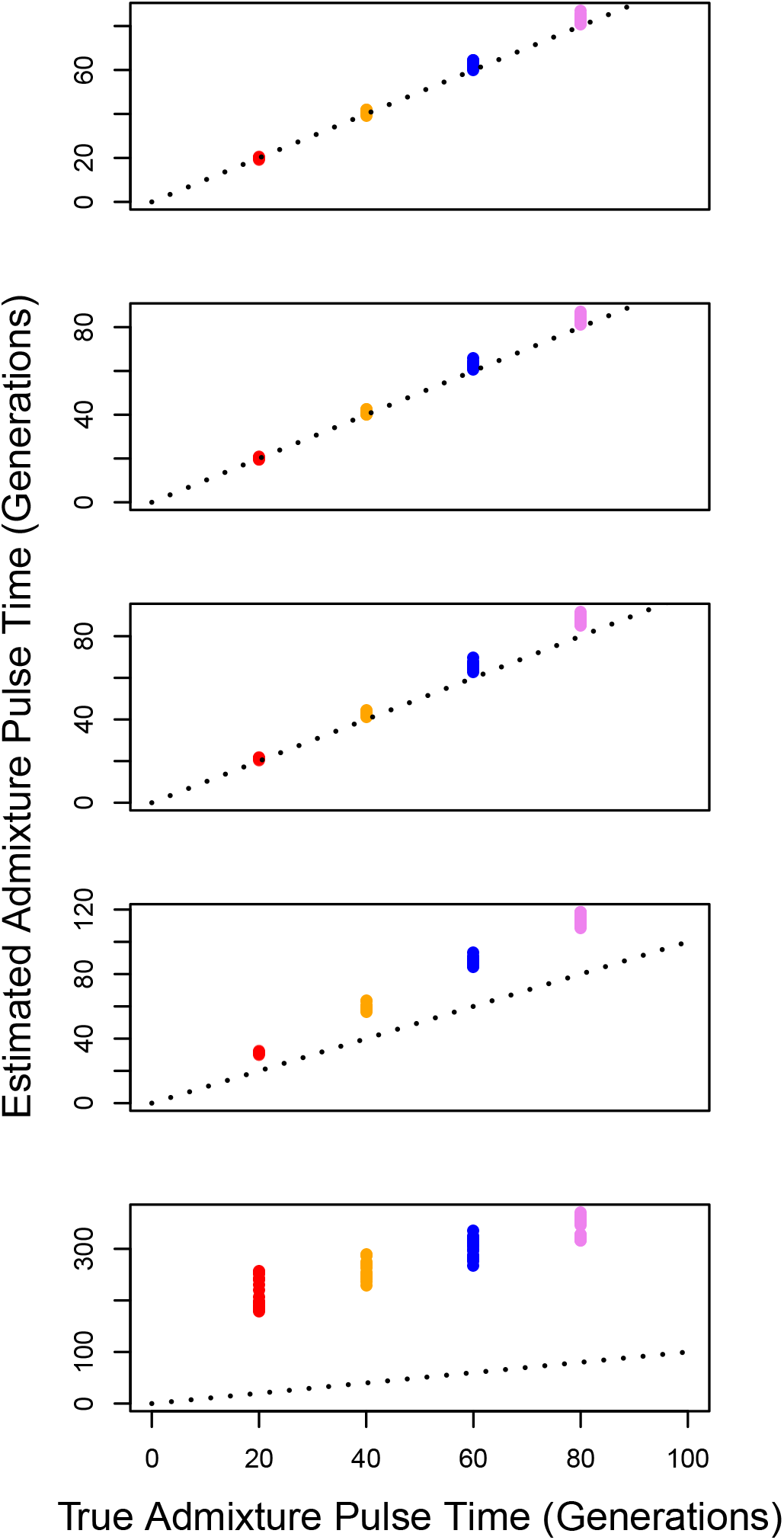
Admixture time estimates for varying divergence between the real source population and assumed reference. Here, we applied a single pulse admixture model to single pulse admixture data. From top to bottom, the reference used to estimate the timing of admixture was 0.005, 0.01, 0.02, 0.05, and 0.1 N_*e*_ generations diverged from the real source population. We simulated single pulse models occurring 20 (red), 40 (orange), 60 (blue), and 80 (pink) generations ago. Note that axes may differ between subplots. Summary statistics of the model fit are listed in Table S8.

**Table S8.** Accuracy statistics of single pulse admixture model fit to single pulse admixture population at varying levels of divergence between source and assumed reference. Listed statistics are described in Table S1.

| divergence | pulse_time | mean_posterior_error | accuracy | confident_correct |
| --- | --- | --- | --- | --- |
| 0.005 | 20 | 0.0002 | 0.9977 | 0.9927 |
| 0.005 | 40 | 0.0005 | 0.9952 | 0.9847 |
| 0.005 | 60 | 0.0008 | 0.9926 | 0.9764 |
| 0.005 | 80 | 0.0011 | 0.9901 | 0.9680 |
| 0.01 | 20 | 0.0003 | 0.9974 | 0.9922 |
| 0.01 | 40 | 0.0006 | 0.9948 | 0.9838 |
| 0.01 | 60 | 0.0009 | 0.9921 | 0.9752 |
| 0.01 | 80 | 0.0012 | 0.9894 | 0.9665 |
| 0.02 | 20 | 0.0003 | 0.9969 | 0.9910 |
| 0.02 | 40 | 0.0007 | 0.9939 | 0.9819 |
| 0.02 | 60 | 0.0011 | 0.9906 | 0.9720 |
| 0.02 | 80 | 0.0015 | 0.9875 | 0.9625 |
| 0.05 | 20 | 0.0012 | 0.9919 | 0.9803 |
| 0.05 | 40 | 0.0020 | 0.9860 | 0.9646 |
| 0.05 | 60 | 0.0028 | 0.9805 | 0.9501 |
| 0.05 | 80 | 0.0034 | 0.9754 | 0.9367 |
| 0.1 | 20 | 0.0158 | 0.9180 | 0.8310 |
| 0.1 | 40 | 0.0172 | 0.9077 | 0.8083 |
| 0.1 | 60 | 0.0182 | 0.8996 | 0.7895 |
| 0.1 | 80 | 0.0189 | 0.8929 | 0.7735 |

**Supplementary Figure S9.**
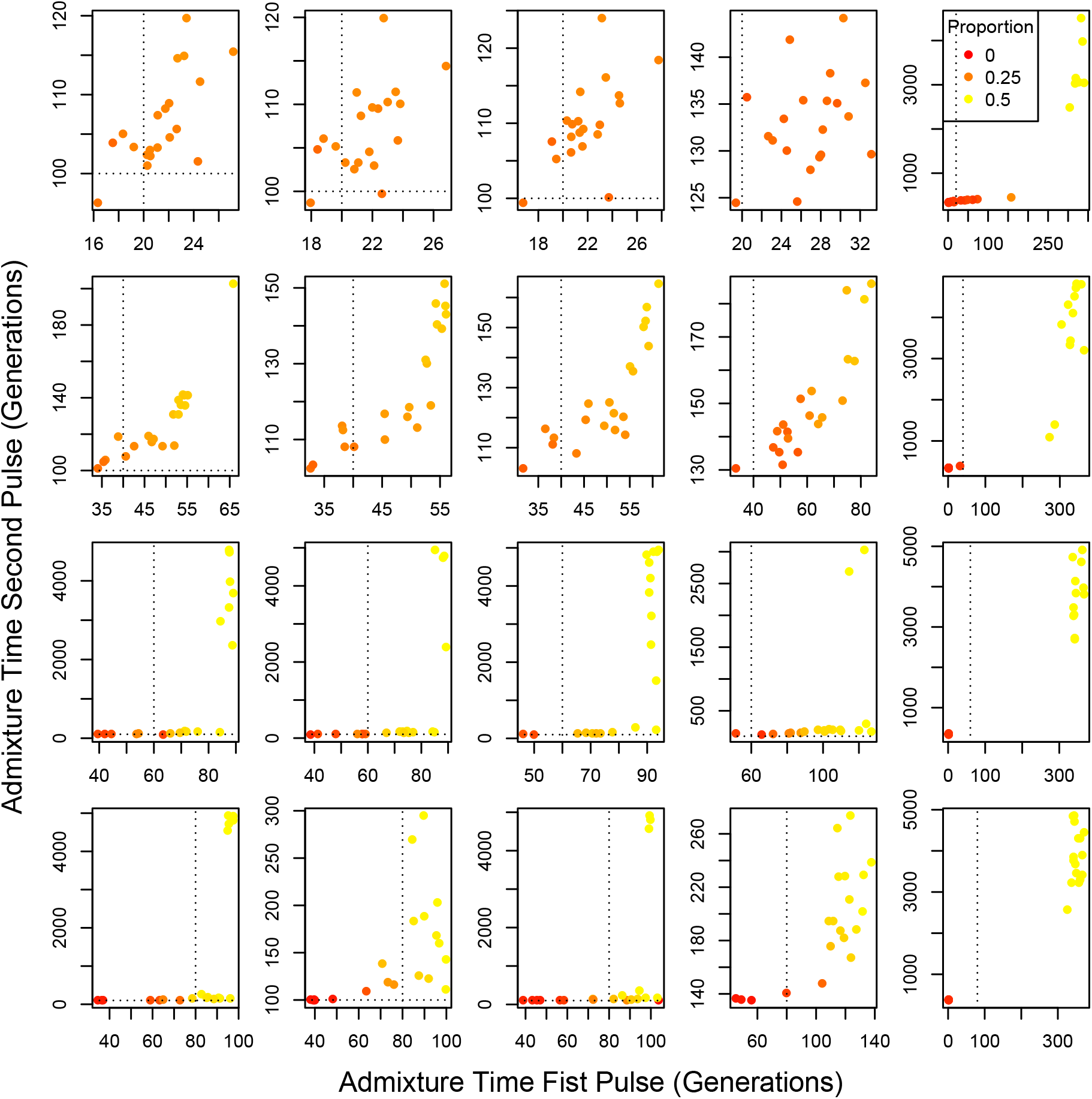
Admixture time estimates for two pulse population model fit to two pulse data at varying level of divergence between source and assumed reference. From left to right, the reference used to estimate the timing of admixture was 0.005, 0.01, 0.02, 0.05, and 0.1 N_*e*_ generations diverged from the real source population. For all admixture models considered, the second pulse occurred 100 generations before the present. From top to bottom, the most recent pulse occurred 20, 40, 60, and 80 generations ago. Note that axes differ slightly between subplots. Summary statistics of model fit are listed in Table S9.

**Supplementary Figure S10.**
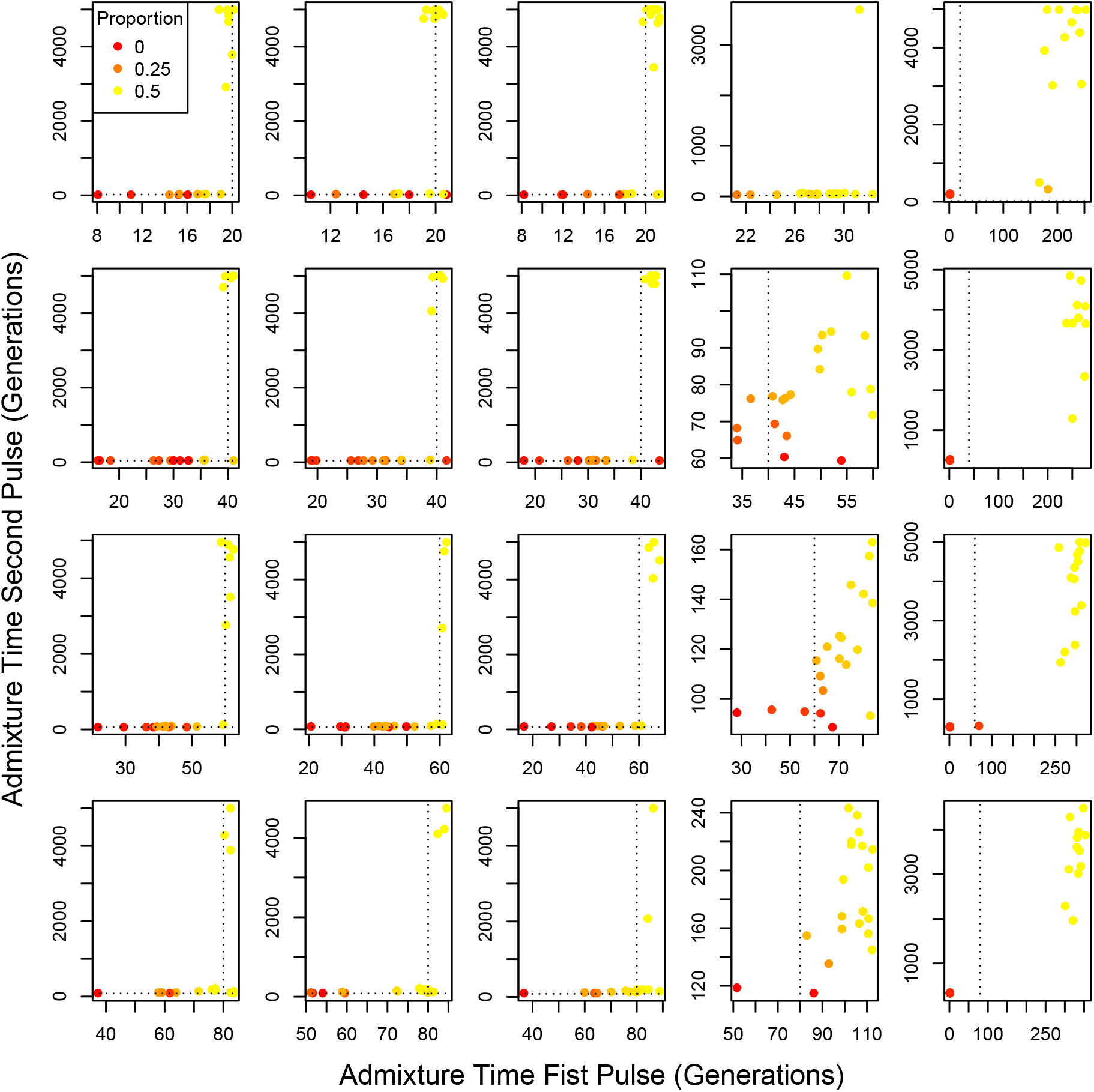
Admixture time estimates for two pulse population model fit to single pulse data at varying level of divergence between source and assumed reference. From left to right, the reference used to estimate the timing of admixture was 0.005, 0.01, 0.02, 0.05, and 0.1 N_*e*_ generations diverged from the real source population. From top to bottom, single pulses occurred 20, 40, 60, and 80 generations ago. Note that axes differ slightly between subplots.

**Table S9.** Accuracy statistics of double pulse admixture model fit to double pulse admixture population at varying levels of divergence between assumed reference and source population. Listed statistics are described in Table S1.

| divergence | pulse_time | mean_posterior_error | accuracy | confident_correct |
| --- | --- | --- | --- | --- |
| 0.005 | 20 | 0.0011 | 0.9910 | 0.9707 |
| 0.005 | 40 | 0.0012 | 0.9902 | 0.9682 |
| 0.005 | 60 | 0.0012 | 0.9893 | 0.9643 |
| 0.005 | 80 | 0.0013 | 0.9883 | 0.9616 |
| 0.01 | 20 | 0.0011 | 0.9903 | 0.9694 |
| 0.01 | 40 | 0.0013 | 0.9894 | 0.9665 |
| 0.01 | 60 | 0.0013 | 0.9885 | 0.9633 |
| 0.01 | 80 | 0.0015 | 0.9875 | 0.9604 |
| 0.02 | 20 | 0.0014 | 0.9887 | 0.9660 |
| 0.02 | 40 | 0.0016 | 0.9876 | 0.9628 |
| 0.02 | 60 | 0.0016 | 0.9865 | 0.9584 |
| 0.02 | 80 | 0.0018 | 0.9853 | 0.9558 |
| 0.05 | 20 | 0.0033 | 0.9772 | 0.9412 |
| 0.05 | 40 | 0.0034 | 0.9755 | 0.9366 |
| 0.05 | 60 | 0.0037 | 0.9739 | 0.9323 |
| 0.05 | 80 | 0.0039 | 0.9721 | 0.9275 |
| 0.1 | 20 | 0.0191 | 0.8904 | 0.7659 |
| 0.1 | 40 | 0.0189 | 0.8900 | 0.7638 |
| 0.1 | 60 | 0.0191 | 0.8887 | 0.7605 |
| 0.1 | 80 | 0.0192 | 0.8854 | 0.7516 |

**Supplementary Figure S11.**
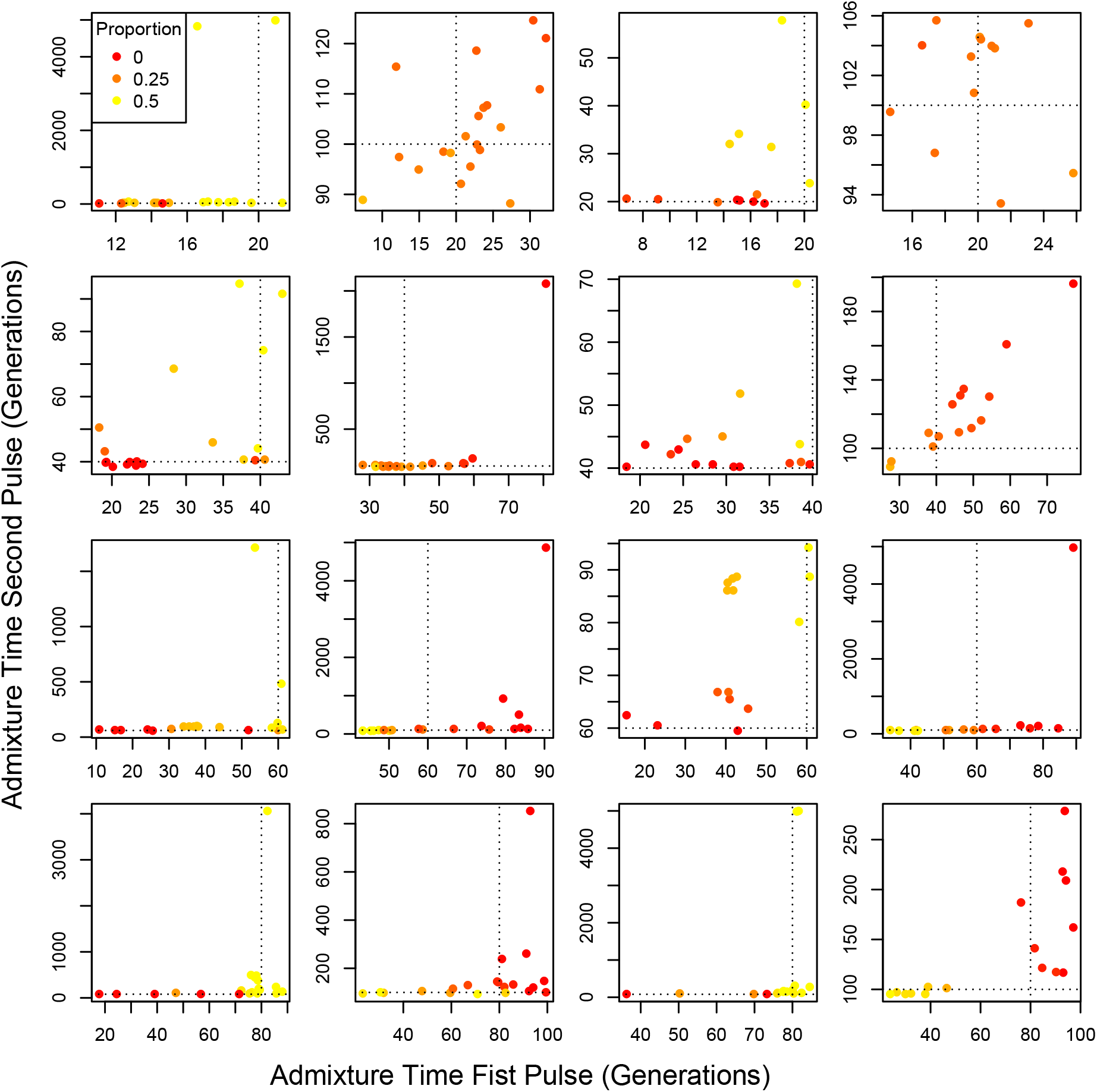
Admixture time estimates for varying sample sizes. From top to bottom, panels show admixture scenarios where the most recent pulse occurred 20, 40, 60, and 80 generations in the past, respectively. For all admixture models considered, the second pulse occurred 100 generations before the present. Columns one and two simulate a sample size of 10 individuals, and columns three and four simulate a sample size of 25 individuals. Columns one and three show simulated single pulse models, columns two and four show simulated double pulse models. In all cases we fit a double pulse model to the simulated data. Summary statistics of model fitting for varying sample sizes are reported in Tables S10, S11, S12. Note that axes may differ between subplots.

**Table S10.** Local ancestry inference accuracy statistics using sample sizes 10 and 25. Listed statistics are described in Table S1.

| admixture_time | sample_size | mean_posterior_error | accuracy | confident_correct |
| --- | --- | --- | --- | --- |
| 20 | 10 | 0.0010 | 0.9911 | 0.9699 |
| 20 | 25 | 0.0010 | 0.9907 | 0.9690 |
| 40 | 10 | 0.0011 | 0.9902 | 0.9668 |
| 40 | 25 | 0.0011 | 0.9900 | 0.9665 |
| 60 | 10 | 0.0012 | 0.9890 | 0.9633 |
| 60 | 25 | 0.0012 | 0.9893 | 0.9639 |
| 80 | 10 | 0.0012 | 0.9883 | 0.9606 |
| 80 | 25 | 0.0013 | 0.9883 | 0.9607 |

**Table S11.** Estimated timing of admixture accuracy statistics using sample sizes 10, 25, and 50. Listed parameters are: simulated timing of admixture pulse (pulse_time), the normalized root mean squared error of estimated admixture time computed from 20 simulations for each sample size tested.

| pulse_time | NRMSE_10 | NRMSE_25 | NRMSE_50 |
| --- | --- | --- | --- |
| 20 | 0.0553 | 0.9634 | 0.0251 |
| 40 | 0.0333 | 0.9806 | 0.0123 |
| 60 | 0.0313 | 0.9902 | 0.0110 |
| 80 | 0.0344 | 0.9723 | 0.0217 |

**Table S12.**
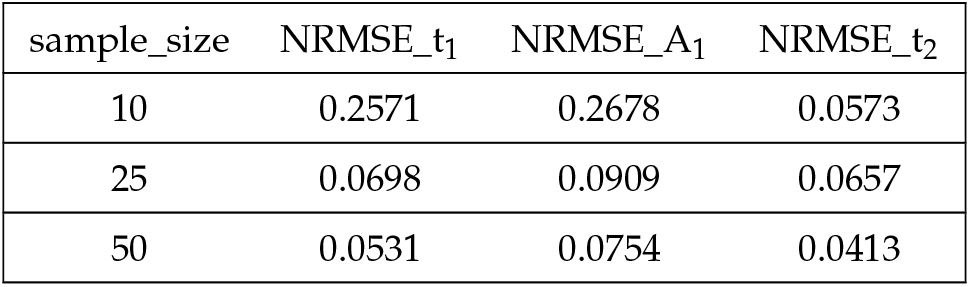
Accuracy statistics of a double pulse model using sample sizes of 10, 25, and 50. Listed statistics are described in Table S1. In addition, we list the normalized root mean squared error of the admixture proportion introduced during *t*_1_ (NRMSE_A_1_).

| sample_size | NRMSE_t<sub>1</sub> | NRMSE_A<sub>1</sub> | NRMSE_t<sub>2</sub> |
| --- | --- | --- | --- |
| 10 | 0.2571 | 0.2678 | 0.0573 |
| 25 | 0.0698 | 0.0909 | 0.0657 |
| 50 | 0.0531 | 0.0754 | 0.0413 |

**Supplementary Figure S12.**
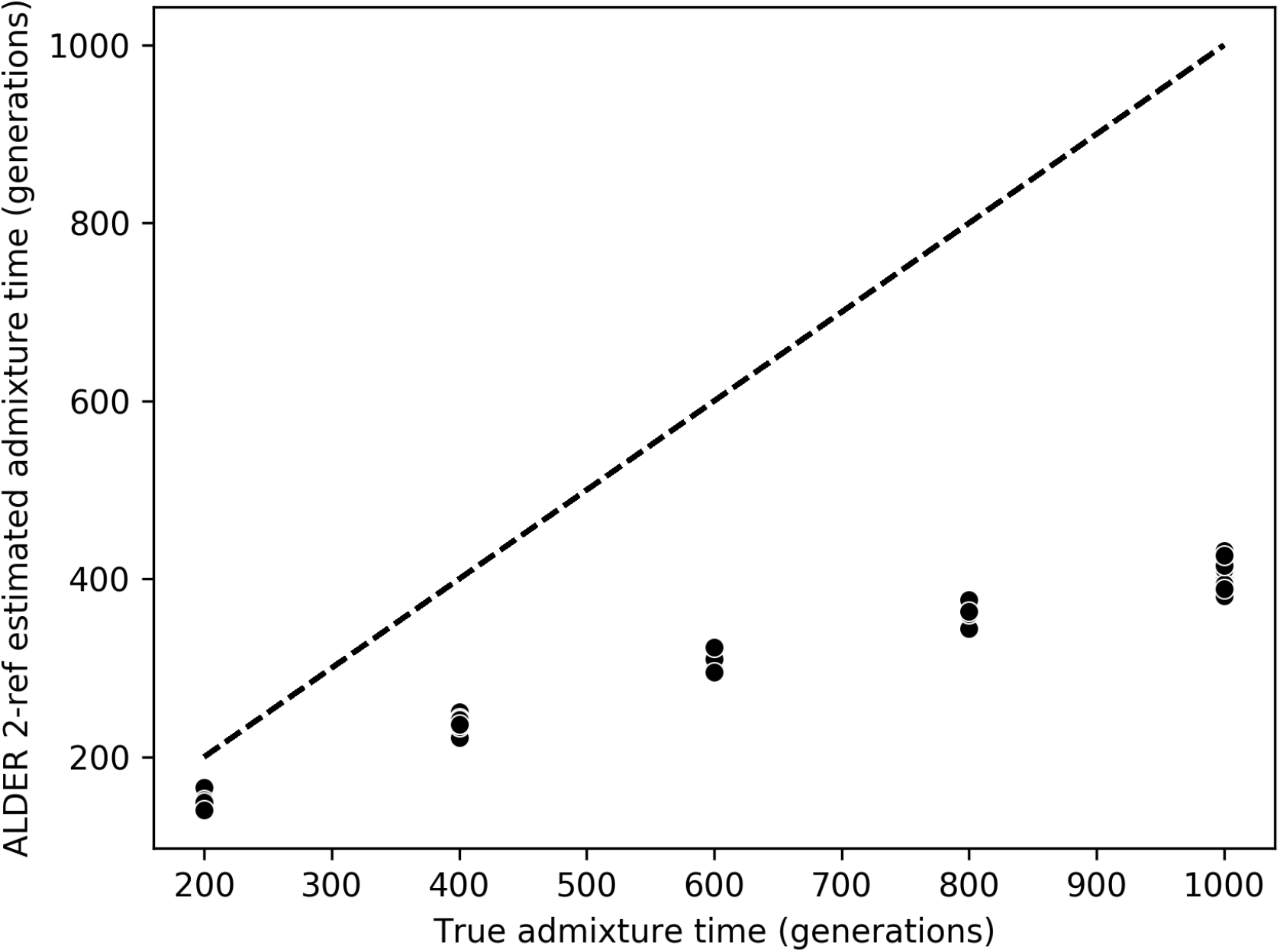
ALDER’s admixture time estimates. We simulated single pulse models occurring 200, 400, 600, 800, and 1000 generations ago (shown left to right). Summary statistics of model fit are listed in Table S14.

**Table S13.** ALDER and Ancestry_HMM NRMSE in estimating timing of single pulse admixture. Listed statistics are described in Table S1.

| pulse_time | ALDER_normalized_NRMSE | Ancestry_HMM_normalized_NRMSE |
| --- | --- | --- |
| 20 | 0.1423 | 0.0251 |
| 40 | 0.1483 | 0.0123 |
| 60 | 0.1777 | 0.0110 |
| 80 | 0.2828 | 0.0217 |
| 100 | 0.2776 | 0.0141 |
| 200 | 0.2581 | 0.0271 |
| 400 | 0.4013 | 0.0234 |
| 600 | 0.4747 | 0.0264 |
| 800 | 0.5362 | 0.0083 |
| 1000 | 0.5933 | 0.0122 |

**Table S14.**
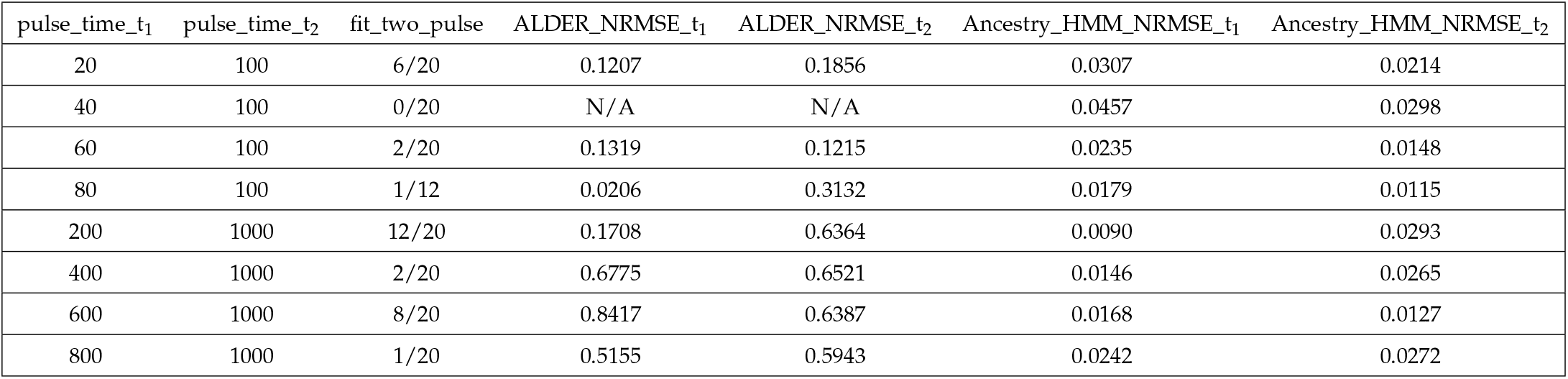
ALDER and Ancestry_HMM NRMSE in estimating timing of double pulse admixture. The table lists the timing of the first admixture pulse in backwards time (pulse_time_t_1_), the timing of the second admixture pulse in backwards time (pulse_time_t_2_), the number of two pulse models fit by ALDER out of 20 replicates (fit_two_pulse), the NRMSE of ALDER’s estimated timing of the first pulse (ALDER_NRMSE_t_1_), the NRMSE of ALDER’s estimated timing of the second pulse (ALDER_NRMSE_t_2_, the NRMSE of Ancestry_HMM’s estimated timing of the first pulse (Ancestry_HMM_NRMSE_t_1_), the NRMSE of Ancestry_HMM’s estimated timing of the second pulse (Ancestry_HMM_NRMSE_t_2_).

